# Phase separation of Mer2 organizes the meiotic loop-axis structure of chromatin during meiosis I

**DOI:** 10.1101/2020.12.15.422856

**Authors:** Bin Tsai, Wei Liu, Dashan Dong, Kebin Shi, Liangyi Chen, Ning Gao

## Abstract

Sexually reproducing organisms acquire genetic diversity through meiotic recombination during meiosis I, which is initiated via programmed DNA double-strand breaks (DSBs) induced by Spo11-containing machinery in each meiotic cell. The combination of programmed DSB sites in each meiotic cell must be diverse, which requires a certain degree of randomness in the distribution of DSBs. The formation of programmed DSBs requires a preestablished loop-axis structure of chromatin. Here, we demonstrate that the axial element protein Mer2 undergoes liquid-liquid phase separation *in vitro* and *in vivo* through its intrinsically disordered C-terminal domain. A DNA binding motif within its central domain is responsible for bringing DNA into Mer2 liquid droplets and Mer2-DNA complex could assemble into filamentous structures extending from the droplets. These results suggest that phase separation of Mer2 drives the formation of a droplet-loop structure of meiotic chromatin to facilitate and to diversify programmed DSB formation.

## Main

Meiosis is the foundation of genetic diversity. Meiotic recombination during prophase of meiosis I recombines the maternal genome and paternal genome to yield a novel genome setup (*1*). Unlike conventional homologous recombination in mitotic cells, meiotic recombination is initiated via programmed DNA double-stand breaks (DSBs) (*2, 3*). Each meiotic cell contains a certain number of programmed DSBs (*4*), and the possible outcome of resulting genetic reshuffling is determined by this specific combination of DSB sites and the choice of subsequent homologous recombination pathways. To achieve genetic diversity, the distribution of DSB sites in each meiotic cell must be different to ensure that each meiotic cell is unique. Theoretically, all sites within the genome are possible for DSB formation, and the simplest way to achieve a maximal number of possible combinations of DSBs is to allow random distribution of DSBs within the entire genome. However, considering the cytotoxicity of DSBs and the physical structure of chromosomes, inappropriate sites of DSBs lead to genome instability; therefore, programmed DSBs are unlikely to be allowed in genomic regions, such as centromeres and repetitive sequences (*5, 6*). Thus, some genomic regions are less likely to contain programmed DSBs compared to other regions, resulting in coldspots and hotspots of programmed DSB formation within a population of meiotic cells. In this regard, programmed DSB formation is random to a certain extent in each meiotic cell and nonrandom within the population of meiotic cells (*7–9*). A central question is to understand the driving force of programmed DSB formation and how it introduces a certain degree of randomness into this process.

Formation of programmed DSBs requires establishment of the loop-axis structure of chromatin (*10, 11*), which arises during prophase of meiosis I. Spo11-associated proteins, including axial element proteins Mer2, Cohesin-Rec8, Hop1-Red1 and Rec114-Mei4, form the proteinaceous axis (*1*). The H3K4 me2/3 reader protein Spp1 binds to chromatin loops and tethers chromatin loops to the axis via its interaction with Mer2 (*12, 13*). Programmed DSB formation mediated by Spo11 occurs in chromatin loops (*14*). To ensure that programmed DSBs are introduced with a certain degree of randomness, loops must be selected with a certain degree of randomness for tethering. Although loops are tethered to the axis via the interaction between Spp1 and Mer2 (*12, 13*), the molecular basis that contributes to this randomness is unclear. In this study, we demonstrate that Mer2 organizes a loop-axis structure together with chromatin DNA and other protein factors via liquid-liquid phase separation, which may introduce randomness to the process of programmed DSB formation.

### Liquid-liquid phase separation of Mer2 in vitro

Liquid-liquid phase separation is responsible for the compartmentalization of cellular contents via the formation of biomolecular condensates, including DNA (chromatin), RNA and protein (*15, 16*). Although phase separation aids in the organization and distribution of biomolecules in most cases, the thermodynamics of liquid condensates should also be sufficient to introduce a certain degree of randomness into biological processes. Multivalent interactions and interactions between intrinsically disordered regions are two common mechanisms for liquid-liquid phase separation (*17*). The central domain of Mer2 is predicted to contain a coiled-coil domain, which is typically involved in multivalent interactions. Additionally, the N- and C-terminal domains of Mer2 as well as its central domain are all predicted to be intrinsically disordered (Fig. 1a, Extended Data Fig. 1) (*18*). This finding prompted us to hypothesize that Mer2 may undergo liquid-liquid phase separation and that the meiotic loop-axis structure of chromatin might actually be organized as a liquid-like biomolecular condensate (Fig. 1a).

**Fig. 1.**
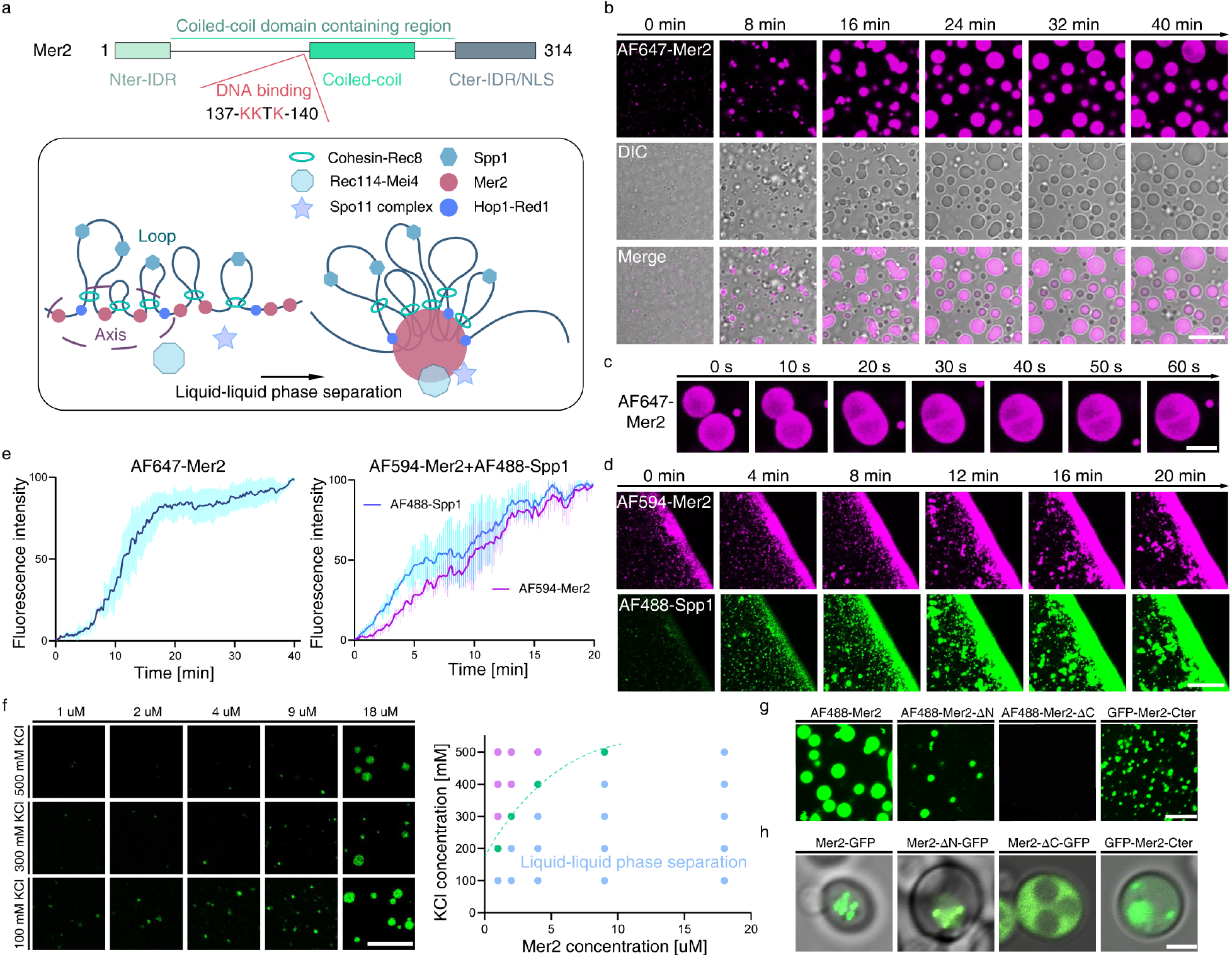
Mer2 undergoes liquid-liquid phase separation via its C-terminal domain. (**a**) Proposed model for organization of the meiotic loop-axis structure via liquid-liquid phase separation of Mer2. (**b**) Time-lapse imaging of Mer2 phase separation. Mer2 (2% was labeled with Alexa Fluor 647, 10 μM) was imaged at room temperature for 40 min. Scale bar, 20 μm. (**c**) Time-lapse micrographs of merging Mer2 droplets. Scale bar, 10 μm. (**d**) Time-lapse imaging of Mer2-Spp1 phase separation. Equal volumes of Mer2 (2% was labeled with Alexa Fluor 594, 10 μM) were mixed with Spp1 (2% was labeled with Alexa Fluor 488, 10 μM) and imaged at room temperature for 20 min. Scale bars, 20 μm. (**e**) Fluorescence intensity of Mer2 phase separation over 40 min and Mer2-Spp1 phase separation over 20 min. Quantification of fluorescence intensity was performed on *n* = 3 fields, and the data shown were normalized to 100% by maximal fluorescence intensity. (**f**) Phase diagram of Mer2 phase separation with varying Mer2 concentrations and KCl concentrations. Scale bar, 10 μm. (**g**) Representative micrographs of liquid droplets formed by wild-type Mer2 and Mer2 mutants. Mer2 variants (2% was labeled with Alexa Fluor 488, 10 μM) and GFP-Cter-Mer2 (25% PEG-3350, 18 μM) were imaged at room temperature. Scale bar, 10 μm. (**h**) Representative micrographs of yeast cells overexpressing Mer2 variants. Scale bar, 2 μm.

To test this possibility, recombinant Mer2 was purified and subjected to an *in vitro* droplet formation assay performed in a buffer of physiological relevance (pH 7.4, 100 mM KCl). The protein solution was stable and clear when incubated on ice. However, it turned turbid quickly after being transferred to room temperature, suggesting a temperature-dependent phase separation of Mer2. This notion is consistent with the fact that temperature affects the efficient progression of meiosis and the landscape of meiotic recombination (*19, 20*). To directly examine the phase separation of Mer2, we used fluorescently labeled Mer2 for the *in vitro* droplet formation assay. Upon transfer to room temperature from ice, fluorescently labeled Mer2 started to form micrometer-sized liquid droplets (Fig. 1b, Supplementary Video 1). We also observed frequent fusion between droplets (Fig. 1c), which led to a gradual increase in the fluorescence intensity in the field of view (Fig. 1e). Next, we plotted the phase diagram of Mer2 phase separation. Of note, increasing the concentration of Mer2 aids in phase separation, whereas increasing ionic strength inhibits phase separation (Fig. 1f, Extended Data Fig.3), suggesting that Mer2 phase separation is driven by interactions among charged residues within IDRs.

Mer2 might serve as a key constituent of a complex biomolecular condensate. Mer2-interacting protein Spp1 colocalizes with Mer2 during meiosis I prophase (*12, 13, 21, 22*). To test whether Spp1 participates in Mer2 phase separation, we incubated fluorescently labeled Spp1 with fluorescently labeled Mer2. Upon mixing, fluorescently labeled Spp1 and Mer2 underwent phase separation together (Fig. 1d, Supplementary Video 2), and Spp1 signals perfectly overlap with those of Mer2 accompanied by increasing fluorescence intensity of both Spp1 and Mer2 over time in the field of view (Fig. 1e). This finding suggests that Spp1 participates in Mer2 phase separation.

To map the domains responsible for liquid-liquid phase separation of Mer2, we constructed Mer2 mutants based on sequence analysis. Mer2 was divided into three parts: the Nter-IDR, Cter-IDR and central coiled-coil containing region. Wild-type Mer2, Mer2-ΔN with the N-terminal domain truncated and Mer2-ΔC with the C-terminal domain truncated were fluorescently labeled and subjected to an *in vitro* droplet formation assay. Wild-type Mer2 and Mer2-ΔN were found to undergo phase separation, whereas no liquid droplet was observed for Mer2-ΔC (Fig. 1g, Supplementary Video 3). We overexpressed these Mer2 variants in yeast cells (tagged with GFP under the control of the CUP1 promoter). Here, wild-type Mer2 and Mer2-ΔN formed bright puncta in yeast nuclei, whereas Mer2-ΔC predominantly localized to the cytoplasm in a dispersed manner (Fig. 1h), suggesting that the C-terminal domain of Mer2 harbors a nuclear localization signal. Furthermore, we purified the C-terminal domain of Mer2 with GFP tagged at its N-terminus and found that the C-terminal domain alone is sufficient to drive phase separation *in vitro* (Fig. 1g). Overexpression of this construct in yeast cells also generated bright puncta in nuclei. These results indicate that Mer2 phase separation is mediated by the C-terminal domain of Mer2.

### Phase separation of Mer2 and DNA

Size-exclusion chromatography of purified full-length Mer2 as well as different domain truncations in high salt buffer (500 mM KCl) at 4°C shows that Mer2 variants self-assembled into large oligomers (Extended Data Fig. 1c-1g). Another observation is that DNA contamination in purified Mer2 samples was high even at such high ionic strength, suggesting that Mer2 might be a DNA binding protein. To test whether Mer2 can bind to DNA *in vitro*, purified Mer2, which is free of DNA contamination, was used for a gel-shift assay. As a result, even in high salt buffer (500 mM KCl), Mer2 exhibited a high affinity to all tested DNA substrates that varied in length and sequence (Fig. 2a, Extended Data Fig. 4e-g).

**Fig. 2.**
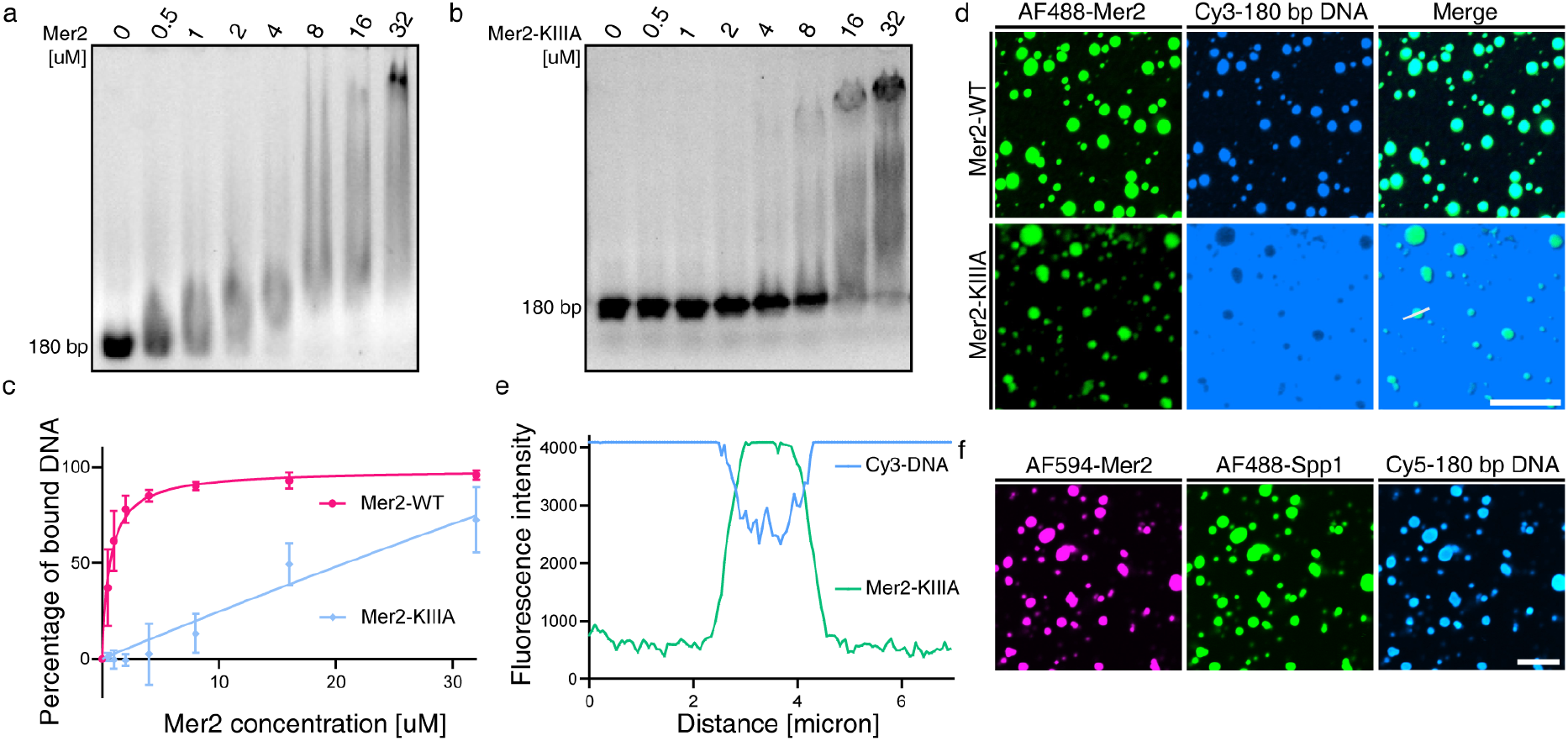
DNA binds to Mer2 and participates in phase separation via interaction with its KKTK motif. (**a**) Gel-shift assay using varying concentrations of wild-type Mer2 to shift 0.1 μM 180 bp DNA in high salt buffer (500 mM KCl). (**b**) Same as (**a**) but for Mer2-KIIIA mutant (K137A, K138A and K140A). (**c**) Quantification of gel-shift assay for wild-type Mer2 and Mer2-KIIIA from *n* = 3 independent experiments; data were fitted with a hyperbola curve. (**d**) Representative micrographs of Mer2-DNA and Mer2-KIIIA-DNA phase separation. Mer2 (2% was labeled with Alexa Fluor 488, 10 μM) or Mer2-KIIIA (2% was labeled with Alexa Fluor 488, 10 μM) was transferred to room temperature, and 5’-Cy3-180 bp DNA (0.27 μM) was added to each sample and imaged. Scale bar 10 μm. (**e**) Quantification of the fluorescence intensities of Mer2-KIIIA and DNA along the line scan indicated by the white line in the merged image of Mer2-KIIIA in (D). (**f**) Representative micrographs of Mer2-Spp1-DNA phase separation. Equal volumes of Mer2 (2% was labeled with Alexa Fluor 594, 10 μM) and Spp1 (2% was labeled with Alexa Fluor 488, 10 μM) were mixed and transferred to room temperature, and 5’ Cy5-180 bp DNA (0.27 μM) was added and imaged. Scale bar, 10 μm.

To map the DNA-interacting domains, Mer2-ΔN, Mer2-ΔC and Mer2-CC were similarly analyzed in a gel-shift assay, which demonstrated that they all exhibited impaired DNA-binding ability (Extended Data Fig. 4a-d). This finding indicates that all three regions contribute to the interaction with DNA. Mer2 contains a basic loop (residues 137-KKTK-140) within its central coiled-coil domain. We suspect that this highly basic KKTK motif might be responsible for DNA binding. A collection of Mer2 mutants was thus derived by introducing point mutations at these positions, including Mer2-K137A, Mer2-K138A and Mer2-KIIIA (K137A, K138A and K140A). All these Mer2 mutants exhibited impaired DNA binding ability, whereas Mer2-KIIIA is affected the most (Fig. 2b-c, Extended Data Fig. 4h-i). Particularly, the triple mutation of Mer2-KIIIA displays even weaker affinity than those domain truncations (Extended Data Fig. 4d). These results suggest that KKTK might be the primary DNA-interacting motif.

Given that Mer2 is a DNA binding protein, we assume that chromatin DNA might also be involved in Mer2 phase separation. To examine whether DNA can participate in Mer2 phase separation, we incubated fluorescently labeled DNA with wild-type Mer2 and Mer2 mutants. The signal of fluorescently labeled DNA is specifically enriched within liquid droplets formed by wild-type Mer2 and Mer2-ΔN but not Mer2-KIIIA (Fig. 2d, Extended Data Fig. 5, Supplementary Videos 4, 5). Further analysis indicates that DNA is excluded from liquid droplets of Mer2-KIIIA (Fig. 2e), confirming that DNA participates in Mer2 phase separation through interaction with the KKTK motif. In addition, Mer2, Spp1 and DNA can form liquid droplets together (Fig. 2f), suggesting that Spp1 and chromatin DNA are also part of the Mer2 condensate during meiosis.

### In vivo phase separation of Mer2 during meiosis

Although many biomolecules were shown to form condensates via liquid-liquid phase separation *in vitro*, it is important to explore their functional relevance *in vivo* under physiological conditions. To avoid possible artifacts from overexpression of plasmid-borne fluorescently labeled Mer2, we constructed a diploid yeast strain with only one allele of Mer2 tagged with GFP at its C terminus for *in vivo* imaging using SR-FACT (super-resolution fluorescence-assisted diffraction computational tomography). SR-FACT combines three-dimensional optical diffraction tomography (ODT) and two-dimensional fluorescence Hessian structured illumination microscopy (Hessian-SIM) (*23, 24*). ODT is a label-free technique that can identify subcellular structures based on differences in their refractive index. Biomolecular condensates formed by phase separation presumably change their refractive index; thus, they can be visualized by ODT as a subcellular structure. We used ODT to observe biomolecular condensates *in vivo* and Hessian-SIM to identify and track biomolecular condensates discovered from ODT.

Mer2 is a meiosis-specific protein, and production of Mer2 protein only starts after initiation of meiosis. Mer2 is synthesized in the cytoplasm at the start of premeiotic S phase and enters the nucleus before meiosis I prophase. To observe Mer2 in the cytoplasm, yeast cells were imaged after being transferred into sporulation medium for 2 hours, which corresponds to premeiotic S phase. Mer2-GFP signals appear as fast-moving puncta in the cytoplasm, and ODT reveals that Mer2-GFP signals indeed colocalize with a subcellular structure (Fig. 3a, Supplementary Video 6). These results suggest that Mer2-GFP is a constituent of this structure and that Mer2 could undergo liquid-liquid phase separation *in vivo*. To observe Mer2 in the nucleus, yeast cells were imaged after sporulation for 4 hours, which corresponds to meiosis I prophase. Mer2-GFP signals appear in the nucleus as multiple foci and are extremely dynamic. Since the refractive index of the yeast nucleus is different from that of the cytoplasm and the fine structure of the nucleus cannot be resolved using ODT, the observation of Mer2 phase separation was exclusively based on Hessian-SIM. Coalescence of two separate Mer2-GFP puncta in the nucleus was observed (Fig. 3b, Extended Data Fig. 6a, Supplementary Videos 7-9), reflecting the liquid property of Mer2-GFP puncta. Additionally, fission of Mer2-GFP puncta was also observed (Fig. 3c, Supplementary Video 8). Fusion of Mer2-GFP puncta could be driven by liquid-liquid phase separation, which may aid the organization of meiotic chromatin and related factors, whereas fission of Mer2 droplets is likely due to other forces that drive chromatin dynamics, which counteracts the phase separation of Mer2 (Fig. 3e). These results imply that Mer2 phase separation *in vivo* occurs in a chromatin context and interplays with chromatin dynamics. Given that meiotic nuclei were too dynamic, we chose to overexpress Mer2 in yeast cells to perform fluorescence recovery after photobleaching (FRAP) analysis to examine whether Mer2 puncta are dynamic structures. After photobleaching, we found that the Mer2-GFP signal indeed reappeared after a short time (Fig. 3e-f, Supplementary Video 10), and FRAP analysis on other Mer2 variants yielded similar results (Extended Data Fig. 6b-f). Altogether, our *in vivo* imaging results demonstrate that Mer2 phase separation occurs in cells and should have an important biological function in meiosis.

**Fig. 3.**
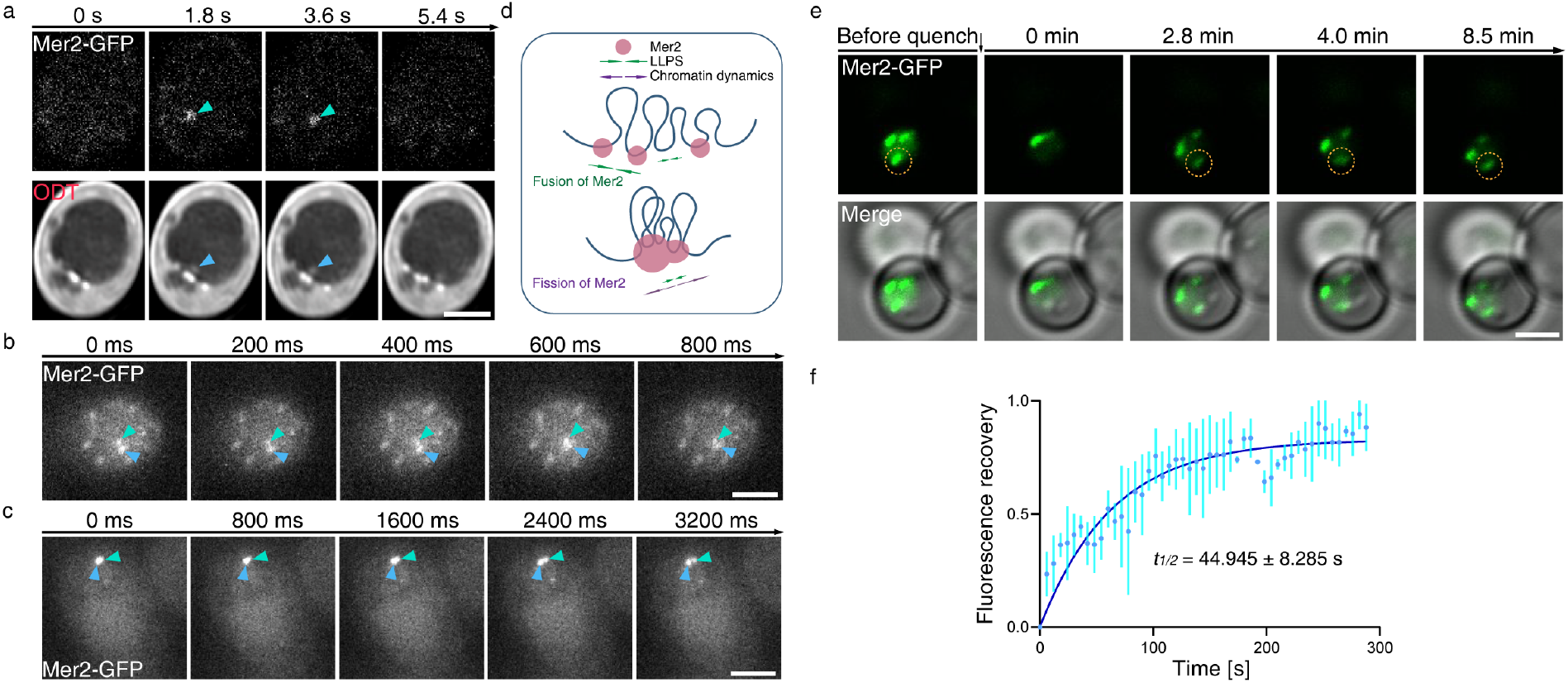
Live-cell imaging of Mer2 phase separation. (**a**) Live-cell imaging of Mer2-GFP puncta formation in the cytoplasm using SR-FACT. Yeast cells were imaged after transfer to sporulation medium for 2 hours. Green arrows indicate Mer2-GFP signals, and blue arrows indicate a subcellular structure with a different refractive index from the surrounding cellular contents. Scale bar, 2 μm. (**b**) Live-cell imaging of fusion of Mer2-GFP puncta in nuclei. Yeast cells were imaged after transfer to sporulation medium for 4 hours. Green and blue arrows indicate two separate Mer2-GFP puncta that are about to fuse. Scale bars, 2 μm. (**c**) Live-cell imaging of fission of Mer2-GFP puncta in nuclei. Green and blue arrows indicate two puncta formed by fission. Scale bar, 2 μm. (**d**) Proposed model explaining fusion and fission of Mer2 puncta. Mer2 is a DNA binding protein that may bind to chromatin DNA *in vivo* and interact with chromatin dynamics. Mer2 phase separation drives fusion of Mer2-GFP puncta, whereas chromatin dynamics that counteract Mer2 phase separation might drive fission of Mer2-GFP puncta. (**e**) Representative micrographs of FRAP analysis of Mer2-GFP before and after quenching. (**f**) Quantification of fluorescence recovery of Mer2-GFP after quenching from *n* = 3 representative experiments (recovery half-time *t*_1/2_ is presented as the mean ± s.d.).

### Mer2-DNA forms a filamentous structure that coexists with Mer2 droplets

Based on the results of *in vitro* and *in vivo* experiments, we conclude that Mer2 likely forms a hub *in vivo* that contains chromatin and Spp1, and other proteins, including Spo11, may participate via direct or indirect interactions with Mer2 or chromatin. We employed electron microscopy (EM) and stimulated emission depletion microscopy (STED) to examine the structural details of Mer2 condensates *in vitro*.

We incubated fluorescently labeled Mer2 with DNA and examined the sample using STED. In contrast to confocal imaging, signals of DNA molecules not only cover Mer2 droplets but also stretch out to form filamentous structures (Fig. 4a, 4b). Notably, these extending filaments are rooted in Mer2 droplets and direct Mer2 to stretch out from droplets.

**Fig. 4.**
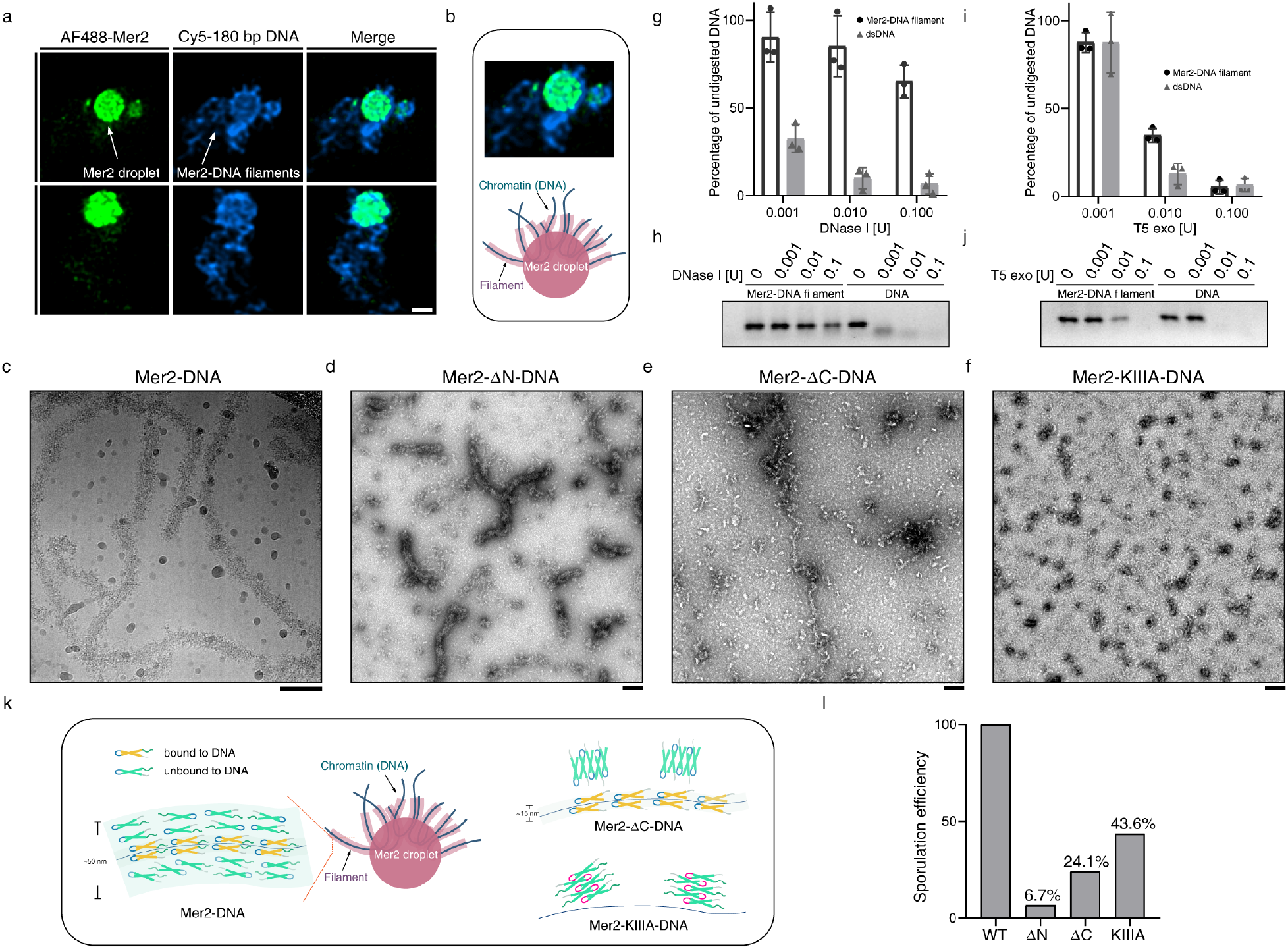
Mer2 forms a filamentous structure with DNA and is organized into a droplet-loop structure. (**a**) STED imaging of Mer2-DNA phase separation. The sample was prepared as described in Fig. 2d. The fluorescence signals of DNA that stretches out from Mer2 droplets represent Mer2-DNA filaments, indicating that the structure observed using STED is a droplet-loop structure that is composed of droplets and filaments. Scale bar, 1 μm. (**b**) Proposed model of droplet-loop structure with Mer2-DNA filaments stretching out from the Mer2 droplet. (**c**) Representative Cryo-EM image of the Mer2-DNA filament. The diameter of the Mer2-DNA filament is not uniform under Cryo-EM, suggesting that formation of the outer layer of the Mer2-DNA filament involves phase separation of Mer2. (**d**), (**e**) and (**f**) Representative negative-staining EM images of Mer2-ΔN-DNA, Mer2-ΔC-DNA and Mer2-KIIIA-DNA. (**g**) Quantification of the DNase I-DNA protection assay from *n* = 3 independent experiments. (**h**) Representative gel image of the DNase I-DNA protection assay. (**i**) Quantification of the T5 exonuclease-DNA protection assay from *n* = 3 independent experiments. (**j**) Representative gel image of the T5 exonuclease-DNA protection assay. (**k**) Proposed model of the formation of the Mer2-DNA filament. (**l**) Sporulation efficiency of yeast strains (SK1 background) carrying Mer2-ΔN, Mer2-ΔC and Mer2-KIIIA mutations. The sporulation efficiency of each strain was calculated from *n* = 3 independent experiments, and the data were normalized to the wild-type strain.

We prepared EM samples of this filamentous structure by titrating Mer2 with DNA. Negative staining EM showed that Mer2 and DNA can form long and flexible filaments (Extended Data Fig. 7a-c) that are micrometers in length. These filaments are much longer than the DNA molecules used, suggesting that they are self-assembled structures. Further cryo-EM indicated that the diameter of these filaments was not uniform (Fig. 4c). Preliminary cryo-EM image processing with these filaments did not reveal any converged structural features, indicating that these filaments were not formed by highly ordered periodic units. This finding also suggests that Mer2 does not simply wrap around DNA with a rigid conformation. We also examined filament formation using Mer2 mutants and DNA (Fig. 4d-f, Extended Data Fig. 7d-f). Mer2-ΔN can form a thick filamentous structure similar to wild-type Mer2; however, the filaments were shorter compared to wild-type Mer2. This finding suggests that the N-terminal domain of Mer2 contributes to axial lengthening of the Mer2-DNA filament (Fig. 4d, Extended Data Fig. 7d). Interestingly, Mer2-ΔC forms thin but long filaments (Fig. 4e, Extended Data Fig. 7e-f). We suspect that this thin filamentous structure is the core of the thick filament formed by wild-type Mer2 and DNA. Additional Mer2 proteins could wrap around the core through phase separation involving intermolecular interactions at the C-terminal domain. Intriguingly, Mer2-KIIIA did not form filamentous structures with DNA and displayed roughly globular structures of typical protein aggregates (Fig. 4f). These results indicate that both properties of DNA interaction and phase separation of Mer2 are required for the formation of long and thick filamentous structures (Fig. 4k).

Given that we were unable to determine the fine structure of the Mer2-DNA filament, we performed a DNA protection assay to determine whether DNA is at the center of the Mer2-DNA filament wrapped up by the Mer2 protein. The Mer2-DNA filament was found to resist DNase I digestion (Fig. 4g-h) but not to T5 exonuclease (Fig. 4i-f-j). These results suggest that DNA is at the center of the Mer2-DNA filament, and Mer2 protects DNA against endonuclease (DNase I) by limiting its accessibility, whereas exonuclease can still digest DNA from the end of the Mer2-DNA filament (Fig. 4k). Furthermore, we measured the sporulation efficiency of yeast strains carrying these Mer2 mutations to assess their effect on meiosis progress and found that Mer2-ΔN, Mer2-ΔC and Mer2-KIIIA all exhibited severely impaired meiosis progress (Fig. 4l), suggesting that DNA binding and phase separation abilities play important roles in meiosis.

### Phase separation of mammalian Mer2 homologs

Sequence analysis suggests that the Mer2 KKTK motif and C-terminal domain of Mer2 could be partially aligned with human and mouse Mer2 homolog IHO1, both of which also contain a coiled-coil domain (Extended Data Fig. 2a-c). We purified two truncated forms of human IHO1 with GFP tagged at their N termini (Extended Data Fig. 2d), GFP-IHO1_1-308_ and GFP-IHO1_255-308_, both of which contain sequences conserved to the C-terminal IDR of Mer2. An *in vitro* droplet assay showed that both could undergo phase separation (Extended Data Fig. 2e). This finding suggests that the ability to undergo phase separation is a conserved function of Mer2 across species.

## Discussion

Taken together, we consider the previously proposed tethered loop-axis model (*10*) to be an incomplete interpretation of a phase separation-based structure, and we designate this phase separation-based structure as a droplet-loop structure (Fig. 4k, Extended Data Fig. 8), which is composed of Mer2 droplets and Mer2-DNA filaments which anchor extended chromatin loops in Mer2 droplets. Mer2 organizes the meiotic chromatin into loop-axis structure via liquid-liquid phase separation to drive programmed DSB formation during meiosis I and offer a certain degree of randomness to the loop-tethering process to diversify programmed DSB formation.

### Phase separation of Mer2 organizes common 3D genome structure and drives programmed DSB formation

Genomes have self-organizing complex 3D structures. The 3D genome structures are diverse among different cell types. A certain type of cell may share a characteristic 3D genome structure with a certain degree of population heterogeneity. Recent advances have revealed several principles of genome organization, including phase separation, chromatin dynamics, polymer-polymer interactions and other principles, such as restriction by nuclear architectural elements (*25*). Dramatic changes in 3D genome structure are observed during the progression of meiosis, and the progress of meiosis is divided into many phases based on shared structural features of the genome of each phase. Meiotic events, such as programmed DSB formation, synapsis and chromosome segregation, all require a characteristic 3D genome structure to proceed (*26*). In the case of programmed DSB formation during prophase of meiosis I, establishment of a loop-axis structure is required (*27*). Our results showed that Mer2 could bind to DNA and undergo phase separation, suggesting that Mer2 could be a driver of 3D genome organization. Mer2 might gain access to axis-associated sequences through other axial element proteins, such as Hop1 and Red1, or simply because Mer2 has free access to these axis-associated sequences at that time point. We propose that Mer2 binds to these axis-associated sequences and undergoes phase separation to organize local loop-axis structures that closely resemble the enhancer-promoter loop structure. Then, these local loop-axis structures condense via phase separation of Mer2 to form a characteristic loop-axis structure that resembles the super-enhancer hub. We designate this structure as a droplet-loop structure (Fig. 1a, Extended Data Fig. 8). Other axis-associated proteins, such as Hop1-Red1, Rec114-Mei4 and the Spo11 complex, might also participate in this process via direct or indirect interactions with Mer2 and chromatin (*10*). The loop-associated protein Spp1 binds to nucleosomes on chromatin loops and then tethers chromatin loops to the axis via phase separation with Mer2 for programmed DSB formation (*12, 13*).

### Phase separation of Mer2 diversifies programmed DSB formation

Given that Spp1 tethers chromatin loops to the axis via phase separation with Mer2, this process will be affected by the thermodynamics of phase separation and local chromatin dynamics (*25*). Within one droplet-loop structure, all chromatin loops are possible for programmed DSB formation. Loops with more Spp1 bound and easy access to Mer2 droplets are more likely to undergo loop tethering and programmed DSB formation, whereas other loops with less Spp1 bound or restricted access to Mer2 droplets are less likely, resulting in loops with much higher priority. Given that only a limited amount of programmed DSBs can be introduced to the genome, a certain set of high priority loops might occupy the chances for programmed DSB formation, limiting the outcome of meiotic recombination. Thus, there must be some mechanisms that could introduce a certain degree of randomness to this process. We propose that the thermodynamics of phase separation and local chromatin dynamics could contribute randomness to this process. In this case, local chromatin dynamics might not alter Spp1 distribution, but the accessibility to Mer2 droplets of each chromatin loop will be affected over time. Additionally, thermodynamics of phase separation might randomly prohibit or aid phase separation of Spp1 with Mer2. As a result of these two effects, although meiotic cells share common 3D genome structure and have similar organization of chromatin loops in general, specific 3D genome structure at each time point will have evident population heterogeneity. The priority of chromatin loops will fluctuate within the population. High priority loops will have a better possibility for programmed DSB formation on average, resulting in hotspots of programmed DSB formation statistically. In contrast, low priority loops will still have a good chance for programmed DSB formation, resulting in coldspots statistically. Thus, all chromatin loops, regardless of their intrinsic property, will have a chance to compete for programmed DSB formation, which is sufficient to diversify the outcome even though this is not an even match.

### Droplet-loop structure could underlie interference among nearby programmed DSB formation sites

Previous studies have demonstrated that a hotspot of programmed DSB formation will reduce the possibility of programmed DSB formation in nearby regions along a chromosome over a distance of tens of kilobases (*28, 29*). It was proposed that hotspots compete for rate-limiting factors that are not free diffusible in the nucleus, resulting in interference among nearby programmed DSB formation sites. Our results agree with this concept; these rate-limiting factors include the limited surface area of the Mer2 droplet, the limited number of Spo11 complexes and other factors. These factors are not freely diffusible as they are restrained by the phase separation of Mer2. Chromatin loops compete for these factors, whereas hotspots remove most of the factors, reducing the possibility of programmed DSB formation in nearby regions.

**Table 1.**
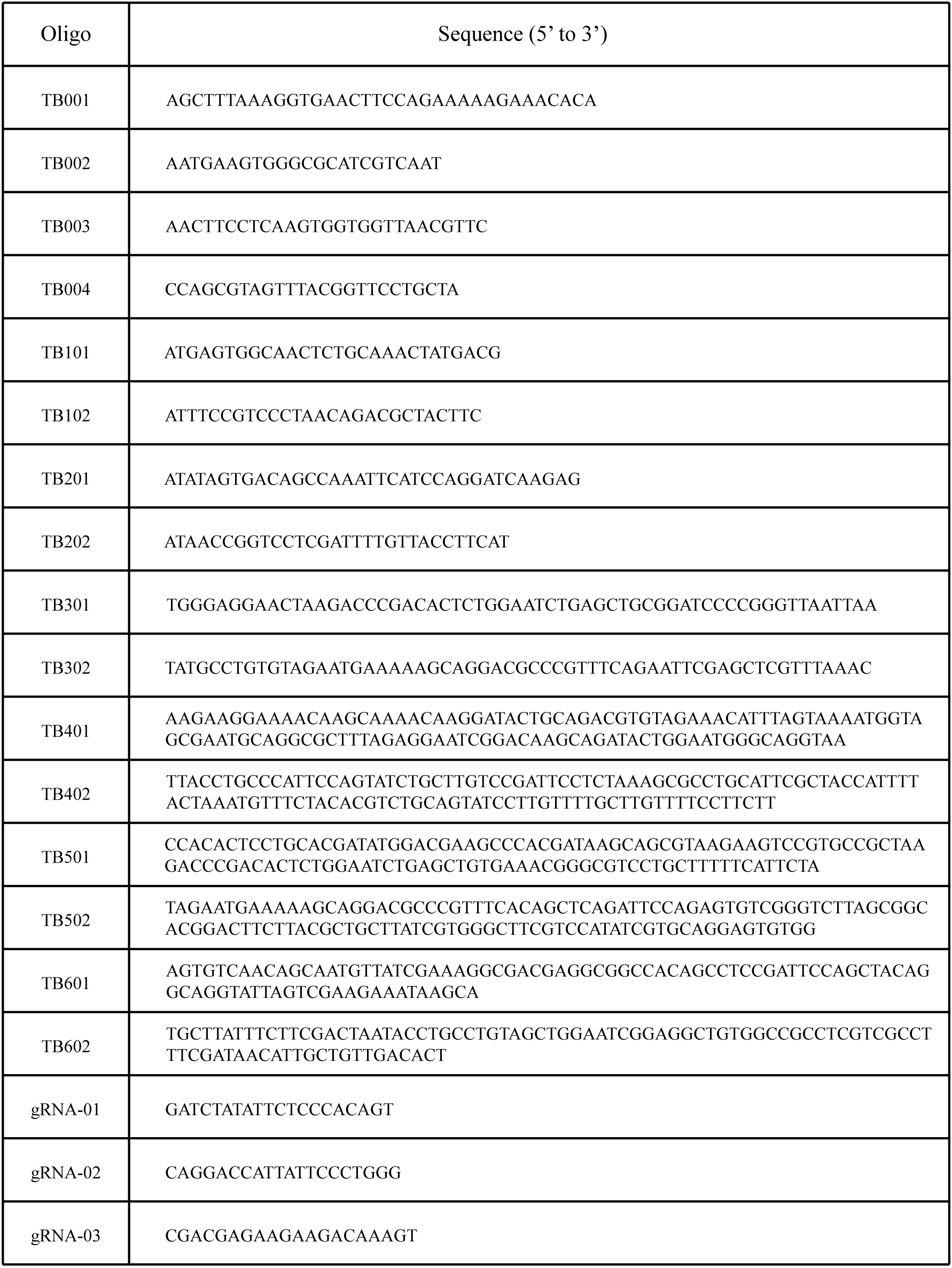
List of oligonucleotides used in this study.

## Acknowledgments

We thank the Core Facilities at School of Life Sciences of Peking University, Electron Microscopy Laboratory, Cryo-EM platform and High-performance Computing Platform of Peking University for assistance with light microscopy, electron microscopy and data processing; Chunyan Shan and Xiaochen Li for help with confocal microscopy and STED microscopy; Yingchu Hu, Yunchao Xie and Pengyuan Dong for help with negative-staining electron microscopy; Liuju Li for help with SR-FACT sample preparation; Damu Wu for help with biochemistry; Yan Chen and Yu Li for help with electron microscopy sample preparation and data analysis; Xuemei Li and Xia Pei for help with Cryo-EM data collection; Jingjing Zhang for help with yeast genetics; Huiqiang Lou for sharing the pFA6a-GFP(S65T)-TRP1 plasmid; and Wei Li for sharing SK1 background yeast strain.

## Funding

The work was supported by the National Key Research and Development Program of China (2019YFA0508904 to N.G.), the National Science Foundation of China (31725007 to N.G.), and the Qidong-SLS Innovation Fund to N.G.

## Author contributions

N.G. and B.T. designed the experiments; B.T. performed biochemistry, electron microscopy, confocal microscopy, yeast genetics and STED microscopy; B.T. constructed a yeast strain for the SR-FACT experiment; W.L. performed SR-FACT and processed SR-FACT data; W.L., D.S.D., K.B.S. and L.Y.C. analyzed SR-FACT data; N.G. and B.T. wrote the manuscript; all authors discussed and commented on the results and the manuscript.

## Competing interests

All authors declare no conflicts of interest.

## Data and materials availability

All data needed to support the conclusions in this study are included in the main text and supplementary materials.

## Supplementary Information

### Materials and Methods

#### Cloning

Genomic DNA of BY4741 background yeast cells was isolated and used for PCR amplification.

For recombinant protein expression and purification, Mer2 variants, IHO1 variants and Spp1 were cloned into *E. coli* expression vectors. To remove the Mer2 intron, exons corresponding to the coding sequences of Mer2 (Mer2WT), Mer2-ΔN (Δ2-42) and Mer2-ΔC (Δ255-304) were amplified by PCR and assembled into a modified pET-22b vector with a C-terminal Strep-Tag II, resulting in pET-22b-Mer2, pET-22b-Mer2-ΔN, and pET-22b-Mer2-ΔC. The coding sequence of Mer2-CC (43-232) was amplified by PCR using pET-22b-Mer2 as a template and assembled into a pET-51b vector with an N-terminal Strep-Tag II, resulting in pET-51b-Mer2-CC. Point mutations were introduced by PCR using pET-22b-Mer2 as a template to yield Mer2-K137A, Mer2-K138A and Mer2-KIIIA (K137A, K1381, K140A), resulting in pET-22b-Mer2-K137A, pET-22b-K138A, and pET-22b-Mer2-KIIIA. For construction of GFP-Mer2-Cter, the coding sequence of GFP was amplified by PCR using pFA6a-GFP(S65T)-TRP1 as a template, the coding sequence of Mer2-Cter (255-314) was amplified by PCR using pET-22b-Mer2 as a template with Strep-Tag II for affinity purification. Then, the two fragments were assembled into the pET-22b vector, resulting in pET-22b-GFP-Mer2-Cter.

IHO1 fragments were amplified by PCR using human cDNA as a template (kindly provided by Jiahua Han), the coding sequence of GFP was amplified by PCR using pFA6a-GFP(S65T)-TRP1 as a template, and IHO1 (1-308) and IHO1 (255-308) were assembled with GFP into a modified pET-51b vector. GFP-IHO1 (1-308) and GFP-IHO1 (255-308) both have Strep-Tag II at their C-terminus.

Spp1 was amplified by PCR and assembled into a modified pET-43.1a vector to yield NusA-tagged Spp1. The original pET-43.1a vector contained a NusA-tag followed by an S-tag and a His-tag, and the His-tag was replaced with a Strep-Tag II for affinity purification, resulting in pET-43.1a-Spp1.

For protein overexpression in yeast cells, Mer2 variants, including Mer2, Mer2-ΔN (Δ2-42), Mer2-ΔC (Δ255-304), Mer2-KIIIA and Mer2-Cter-GFP (255-314), were cloned into a modified pYES2 vector (Thermo Fisher), and the GAL promoter was replaced with the CUP1 promoter for copper-induced protein production. GFP (S65T) was amplified by PCR using pFA6a-GFP(S65T)-TRP1 as a template and fused to the C terminus of each Mer2 variant for fluorescence imaging. The resulting vectors are designated pCUP1-Mer2-GFP, pCUP1-Mer2-ΔN-GFP, pCUP1-Mer2-ΔC-GFP, pCUP1-Mer2-KIIIA-GFP and pCUP1-Mer2-Cter-GFP.

#### Yeast strain construction

A standard PCR-based gene targeting method was used to generate a yeast strain with only one allele of Mer2 tagged with GFP at its C terminus. Primers TB301/TB302 were used to amplify a fragment used for gene targeting from pFA6a-GFP(S65T)-TRP1, and the PCR product was gel purified and used to transform SK1 background yeast cells using the standard LiOAc/ssDNA method. Yeast cells were selected on SD-TRP plates and confirmed by colony PCR and Sanger sequencing.

The CRISPR/Cas9 system was used to generate yeast strains carrying Mer2-ΔN, Mer2-ΔC, and Mer2-KIIIA mutations. A 2 μ vector containing both Cas9 protein and gRNA was constructed for genetic manipulation of yeast. The coding sequence of SpCas9 was amplified by PCR from pX330, and the TEF1 promoter and SNR52 promoter were amplified from genomic DNA of BY4741 background yeast cells. The gRNA scaffold fused to the SUP4 terminator was synthesized, and the vector backbone was amplified by PCR from pYES2 (Thermo Fisher). These fragments were assembled to generate a vector designated pTsGEN-URA. The vector contained the SNR52p-gRNA-SUP4t cassette and TEF1p-SpCas9-CYC1t cassette. The URA3 cassette was replaced with TRP1 and HIS3 to generate pTsGEN-TRP and pTsGEN-HIS, respectively.

For construction of the SK1 background strain carrying the Mer2-ΔN mutation, yeast cells were transformed with pTsGEN-TRP carrying gRNA-01 and repair template, and the repair template was prepared by annealing primers TB401/402. Yeast cells were selected on SD-TRP plates and confirmed by colony PCR and Sanger sequencing.

For construction of the SK1 background strain carrying Mer2-ΔC and Mer2-KIIIA, the same method described for the Mer2-ΔN strain was used with the exception that the gRNA for Mer2-ΔC was gRNA-02 and the repair template was TB501/TB502, the gRNA for Mer2-KIIIA was gRNA-03 and the repair template was TB601/TB602.

#### Protein expression and purification

All proteins were expressed and purified from *Escherichia coli*. For expression of Mer2, Mer2 variants and IHO1 variants, *E. coli* strain Transetta (TransGen) was transformed with plasmids encoding each of the Mer2 variants. *E. coli* cells were induced for protein expression with 0.5 mM IPTG at 16°C overnight. *E. coli* cells were collected by centrifugation. All procedures were performed either on ice or at 4°C. *E. coli* cells were lysed by sonication on ice in buffer H (50 mM Tris-HCl, pH 7.4, 500 mM KCl, 10% glycerol, 0.01% Triton X-100) with 1 mM PMSF and protease inhibitor cocktail (Mei5bio). After sonication, the cell lysate was centrifuged at 30,000 x g, and the supernatant was collected. Streptactin beads 4FF (Smart-Lifesciences) were equilibrated with buffer H and incubated with supernatant for 20 min, washed extensively with buffer H, and eluted with buffer H/2.5 mM d-desthiobiotin (Sigma). Eluted protein was subjected to size-exclusion chromatography in buffer H using a Superose 6 increase 10/300 GL (GE Healthcare) column, and peak fractions with A260 absorbance less than A280 absorbance were considered to be free of DNA contamination. Fractions collected were concentrated by ultrafiltration in buffer H and flash frozen in liquid nitrogen for storage at −80°C.

NusA-tagged Spp1 was expressed and purified in *E. coli* using the same method described for Mer2 with the exception that Spp1 expression was induced in the presence of 0.3 mM IPTG at 16°C overnight. To ensure the stability of the Spp1 protein, the NusA-tag was not removed for subsequent experiments.

#### Fluorophore labeling of proteins

Purified Mer2 was labeled with Alexa Fluor™ 488, Alexa Fluor™ 594 and Alexa Fluor™ 647 (Thermo Fisher). Purified Mer2 protein was buffer exchanged into buffer F (50 mM HEPES-KOH, pH 8.2, 500 mM KCl, 10% glycerol, 0.01% Triton X-100) by ultrafiltration and concentrated to 1 mg/ml. NHS ester of each fluorophore was dissolved with DMSO to 10 μg/μl, and 1 μl of each fluorophore was added into 100 μl of protein solution. Each sample was protected from light and incubated at room temperature on a rotational mixer for 1 hour. Fluorophore labeling was ended by diluting each sample into 10 ml buffer H followed by ultrafiltration. Fluorophore-labeled protein samples were flash frozen in liquid nitrogen in small aliquots for storage at −80°C. Using the method described above, purified Mer2 mutants, including Mer2-ΔN, Mer2-ΔC and Mer2-KIIIA, were labeled with Alexa Fluor™ 488, and purified NusA-tagged Spp1 was also labeled with Alexa Fluor™ 488.

#### DNA substrates and fluorophore labeling of DNA

All primers, including 5’-Cy3 and 5’-Cy5 modified primers, were purchased from Ruibiotech. Primers were used directly for PCR using genomic DNA of BY4741 background yeast cells as template. Here, 90 bp DNA was amplified with TB001/TB002, 180 bp DNA with TB001/TB003, 600 bp DNA with TB201/TB202, and 3051 bp DNA with TB001/TB004. To generate fluorescently labeled DNA, 5’-Cy3- and 5’-Cy5-modified TB001 was used for the preparation of Cy3-180 bp DNA, Cy5-180 bp DNA and Cy5 3051 bp DNA. The PCR mixture for each DNA substrate was subjected to EtOH precipitation at −80°C overnight. DNA was pelleted by centrifugation at 4°C, washed with 70% EtOH and air dried for storage at −40°C.

#### Gel-shift assay

Purified proteins were incubated with corresponding DNA substrates on ice for 10 min in buffer H. DNA loading buffer was then added to each sample, and reaction mixtures were loaded onto a 0.8% agarose gel prepared with TBE buffer. Horizontal electrophoresis cells were placed on ice, reaction mixtures were resolved in ice cold 1x TBE buffer, and DNA substrates were visualized by SYBR™ Gold Nucleic Acid Gel Stain (ThermoFisher). Quantification of gel-shift assays was performed using FIJI (NIH) following the standard method.

#### Negative-staining electron microscopy and cryo-electron microscopy

For negative-staining electron microscopy, wild-type Mer2 and Mer2 mutants were titrated with DNA in buffer L (50 mM Tris-HCl, pH 7.4, 100 mM KCl, 10% glycerol, 0.01% Triton X-100). EM samples were prepared by mixing Mer2 variants with 180 bp DNA at a molar ratio of 20:1 unless otherwise indicated. For the Mer2-DNA filament using 3051 bp DNA, the EM sample was prepared with an equal amount of weight of 3051bp DNA. Carbon-coated copper grids (Zhong Jing Ke Yi) were glow-discharged for 30 s using a plasma cleaner (Harrick PDC-32G-2). Four μl of each sample was applied to a glow-discharged grid and incubated for 1 min, and the excessive solution was removed using filter paper. Samples were stained with 2% uranyl acetate. EM grids were examined under a Tecnai G2 20 Twin microscope (FEI) operated at 120 kV, and images were acquired using a 4k x 4k CCD camera (Eagle, FEI).

For cryo-electron microscopy, Mer2 was mixed with 180 bp DNA at a molar ratio of 20:1. Cryo grids were prepared using Vitrobot Mark IV (FEI) at 4°C and 100% humidity. Four microliters of sample were applied to glow-discharged holey carbon copper grids (Quantifoil, R 1.2/1.3), blotted with filter paper and plunged into liquid ethane. A Titan Krios G3 (FEI) equipped with a K2 summit direct electron detector (Gatan) was used for image collection.

#### Live-cell imaging of yeast cells overexpressing Mer2 variants and fluorescence recovery after photobleaching analysis

Yeast cells of the BY4741 background were transformed with pCUP1-Mer2 vectors and selected on SD-URA plates at 30°C. Selected clones were cultured in SD-URA medium at 30°C. For copper-induced protein production, 0.1 mM CuSO_4_ was added to the medium and cultured for 1 hour at 30°C. Yeast cells were then collected by centrifugation and resuspended in fresh SD-URA medium supplemented with 0.1 mM CuSO_4_ at the proper dilution. Yeast cells were transferred to a glass bottom dish for confocal imaging using a Nikon A1R+ confocal microscope with a 100x oil immersion objective equipped with a 488-nm laser.

For fluorescence recovery after photobleaching analysis using a 488-nm laser, the laser power was set to 80% to bleach a 0.2 μm × 0.2 μm square region of interest. The GFP signal was quenched by bleaching with a 488-nm laser for 1 s, and subsequent time-lapse images were acquired using the A1 stimulation tool and ND stimulation tool in NIS-elements with 6 s intervals while the laser power was set to 8%. Quantification of fluorescence recovery after photobleaching was performed using FIJI (NIH) following the standard method.

#### Confocal fluorescence microscopy and *in vitro* phase separation assay

All samples for the *in vitro* droplet formation assay were imaged using a Nikon A1R+ confocal microscope with a 60x oil immersion objective equipped with 488-nm, 561-nm and 640-nm lasers. Time-lapse images were captured using the ND acquisition tool in NIS elements. Samples were imaged in glass bottom dishes. All glass bottom dishes used were coated with 20 mg/ml BSA for 10 min at room temperature. In general, all fluorescently labeled proteins in buffer H were diluted with corresponding unlabeled proteins to achieve a mixture with 2% protein molecules labeled with fluorophore.

For *in vitro* phase separation assays with only Mer2 protein including AF647 labeled Mer2 along with AF488 labeled Mer2-ΔN, Mer2-ΔC and Mer2-KIIIA, protein samples were stored in buffer H on ice. The salt concentration was adjusted to 100 mM KCl with buffer N (50 mM Tris-HCl, pH 7.4, 10% glycerol, 0.01% Triton X-100), diluted to 10 μM using buffer L on ice and transferred to a glass bottom dish at 19°C for imaging.

For assays using IHO1 variants, the salt concentration was adjusted to 100 mM KCl with buffer N and 50% PEG-3350 solution, diluted with buffer L on ice and transferred to a glass bottom dish at 19°C for imaging.

For assays using Mer2 variants and DNA, AF488-labeled Mer2, Mer2-ΔN and Mer2-KIIIA protein samples were adjusted with buffer N, diluted to 10 μM with buffer L on ice, and transferred to a glass bottom dish at 19°C for imaging. Fluorescently labeled DNA was added to each sample after 20 min and imaged.

For the assay using Mer2 and Spp1, AF594-labeled Mer2 and AF488-labeled NusA-tagged Spp1 were adjusted with buffer N and diluted to 10 μM with buffer L on ice. 10 μl of Mer2 was transferred to a glass bottom dish at 19°C, and 10 μl of Spp1 was added dropwise into Mer2 after 5 min and imaged for an additional 20 min.

For the assay using Mer2, Spp1 and DNA, AF594-labeled Mer2 and AF488-labeled NusA-tagged Spp1 were adjusted with buffer N and diluted to 10 μM with buffer L on ice, and equal volumes of Mer2 and Spp1 were mixed and transferred to a glass bottom dish at 19°C. Fluorescently labeled DNA was added after 20 min and imaged.

#### Super-resolution fluorescence-assisted diffraction computational tomography (SR-FACT) and *in vivo* phase separation assay

SR-FACT consists of a label-free three-dimensional optical diffraction tomography (ODT) module and a two-dimensional fluorescence Hessian structured illumination microscopy module (*23, 24*). The 3D refractive index value distribution, which represents different subcellular structures, can be reconstructed by the ODT module based on wide-field digital holograms and tomographic illumination. The ODT module achieves a lateral resolution of 200 nm and at a volumetric imaging speed of 0.8 Hz (40 × 40 × 20 μm^3^) when a 561-nm single longitudinal-mode laser was used as the illumination source. To eliminate the aberration introduced by the coverslip, extra ODT data of each coverslip at empty areas were taken for calibration of imaging reconstruction. In the Hessian 2D-SIM module, a 488-nm single longitudinal-mode laser is used as the illumination source, which achieves a spatial resolution of 88 nm and imaging at one order of magnitude lower than the photon dose used by conventional SIM. The 2D-SIM was used to guide the interpretation of structures observed by the ODT module. Compared with previous methods (*30–32*), the new dual-mode imaging microscopy technique has great advantages when used for time-lapse correlated super-resolution imaging due to its high spatiotemporal resolution of label-free live cell imaging. Detailed experimental settings for SR-FACT data collection and processing were the same as described previously (*23*).

Yeast cells of the SK1 strain background were used for SR-FACT imaging. The GFP (S65T) coding sequencing with a TRP1 expression cassette was amplified from the pFA6a-GFP(S65T)-TRP1 vector and used for C-terminal tagging of Mer2 at its native locus using a standard PCR-based method, and genotyping confirmed one positive clone with only one allele tagged with GFP. GFP-tagged yeast cells were inoculated into YPD medium, and overnight cultures were diluted in YPA medium (1% yeast extract, 2% peptone, 2% potassium acetate) to OD600 = 0.3 and cultured at 30°C for l0-12 hours. Yeast cells were collected by centrifugation and washed thrice with sporulation medium (2% potassium acetate). Cell pellets were resuspended in sporulation medium to OD600 = 1.9 and cultured at 30°C. For imaging using SR-FACT, yeast cells were collected and diluted with sporulation medium at each time point and imaged in a glass bottom imaging chamber at 23°C.

#### DNA protection assay

Mer2-DNA filaments were prepared using the same method described for electron microscopy experiments. Mer2-DNA filaments that corresponded to 38 ng DNA and naked DNA of the same amount were used as substrates for each experiment.

For the DNase I protection assay, Mer2-DNA filaments and naked DNA were mixed with the corresponding amount of DNase I (NEB) in DNase I reaction buffer (NEB) on ice, incubated at 37°C for 15 min and inactivated at 70°C for 5 min using a thermocycler. DNA loading buffer was added, and the samples were loaded onto a 0.8% agarose gel prepared with TBE buffer. Reaction mixtures were resolved in ice cold 1x TBE buffer.

For the T5 exonuclease-protection assay, Mer2-DNA filaments and naked DNA were mixed with the corresponding amount of T5 exonuclease (NEB) in NEBuffer 4 (NEB) on ice and incubated at 37°C for 15 min followed by 70°C inactivation for 5 min using a thermocycler. DNA loading buffer was added, and the samples were loaded onto a 0.8% agarose gel prepared with TBE buffer. Reaction mixtures were resolved in ice cold 1x TBE buffer.

Quantification of DNA protection assays was performed using FIJI (NIH) following the standard method.

#### Sporulation efficiency analysis

Yeast strains of the SK1 background carrying Mer2-ΔN, Mer2-ΔC and Mer2-KIIIA together with the wild-type SK1 strain were cultured in YPD medium at 30°C for 24 hours. Yeast cells were collected and washed thrice with YPA medium (1% yeast extract, 2% peptone, 2% potassium acetate), resuspended in YPA medium to OD600 = 0.3 and cultured at 30°C for an additional 10 hours. Yeast cells were collected and washed thrice with prewarmed sporulation medium (2% potassium acetate), resuspended in sporulation medium to OD600 = 1.9 and cultured at 30°C for 11 hours. Yeast cells were then collected and resuspended in fresh sporulation medium and transferred to a glass bottom dish for confocal imaging. Yeast cells were imaged using a Nikon A1R+ confocal microscope with a 100x oil immersion objective equipped with a 488-nm laser.

A total of 1,000 cells were counted for each strain from 3 biological replicates, and the percentage of yeast tetrad was calculated. The percentage of yeast tetrads for each strain was normalized to the wild-type SK1 strain to represent sporulation efficiency.

#### Stimulated emission depletion microscopy

For observation of the droplet-loop structure, AF488-labeled Mer2 in buffer H was adjusted with buffer N, diluted to 10 μM with buffer L, and transferred to a glass bottom dish at room temperature. Cy5-180 bp DNA was added after 10 min. The sample was imaged using a Leica TCS SP8 STED 3X microscope with a 100x oil immersion objective equipped with a white light laser (470-670 nm) for excitation. Here, 592-nm and 775-nm lasers were used as depletion lasers. Images were acquired using LAS X Smart STED wizard, and deconvolution of images was performed using Huygens Software.

---

**Extended Data Fig. 1.**
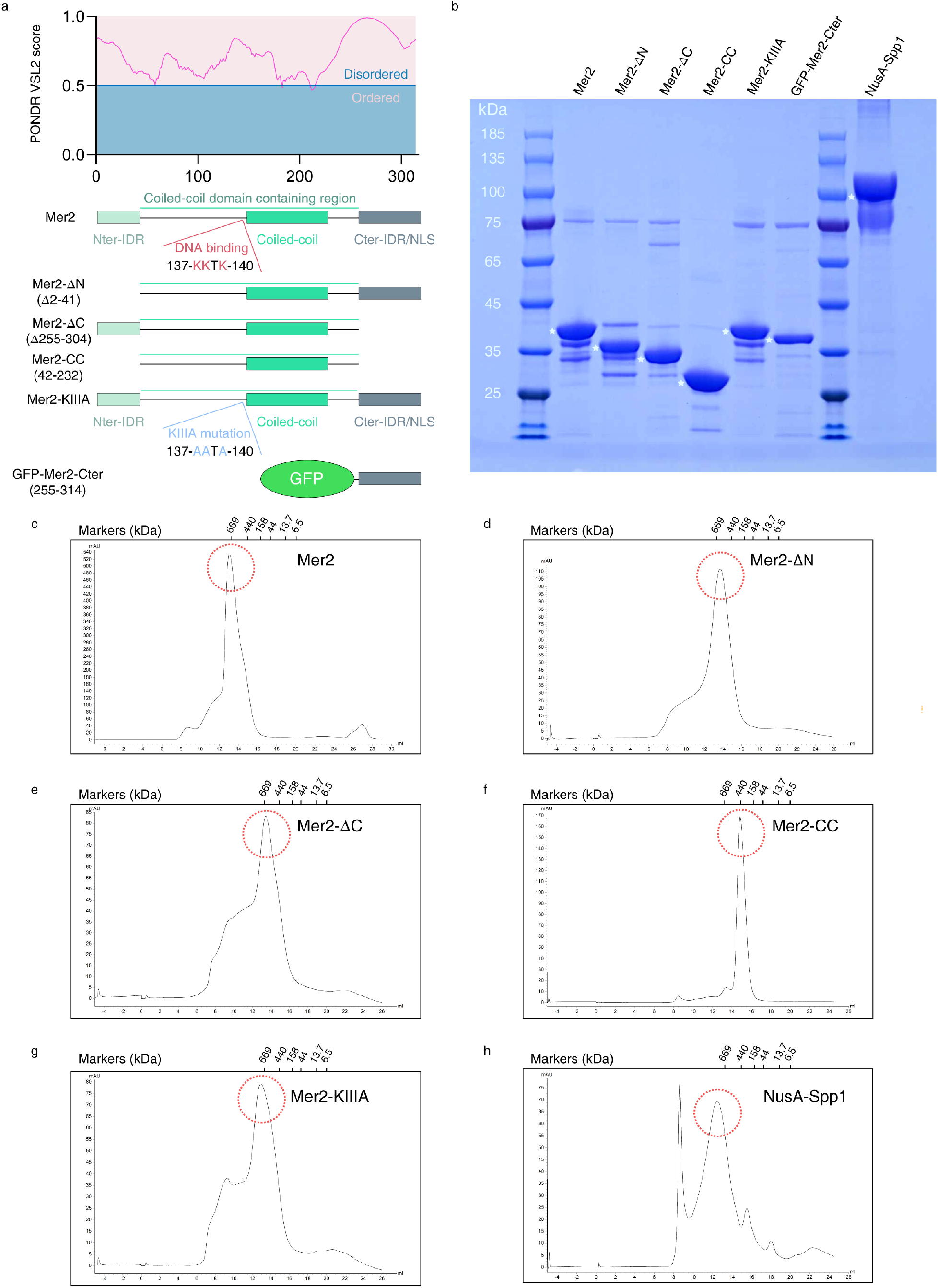
Purification of recombinant Mer2 variants and Spp1. (**a**) Prediction of naturally disordered regions of Mer2 using VSL2 and the domain organization of Mer2 variants. (**b**) SDS-PGAE of Mer2 variants and NusA-tagged Spp1. Approximately 5 μg of each protein was loaded onto a 4-12% SurePAGE gradient gel (Genscript) running in MOPS/SDS buffer and stained with Coomassie brilliant blue. Asterisks indicate proteins of interest. (**d**) to (**h**) Size-exclusion chromatography of Mer2 variants and NusA-tagged Spp1 in high salt buffer (pH 7.4, 500 mM KCl) at 4°C using Superose 6 increase, 10/300 GL.

**Extended Data Fig. 2.**
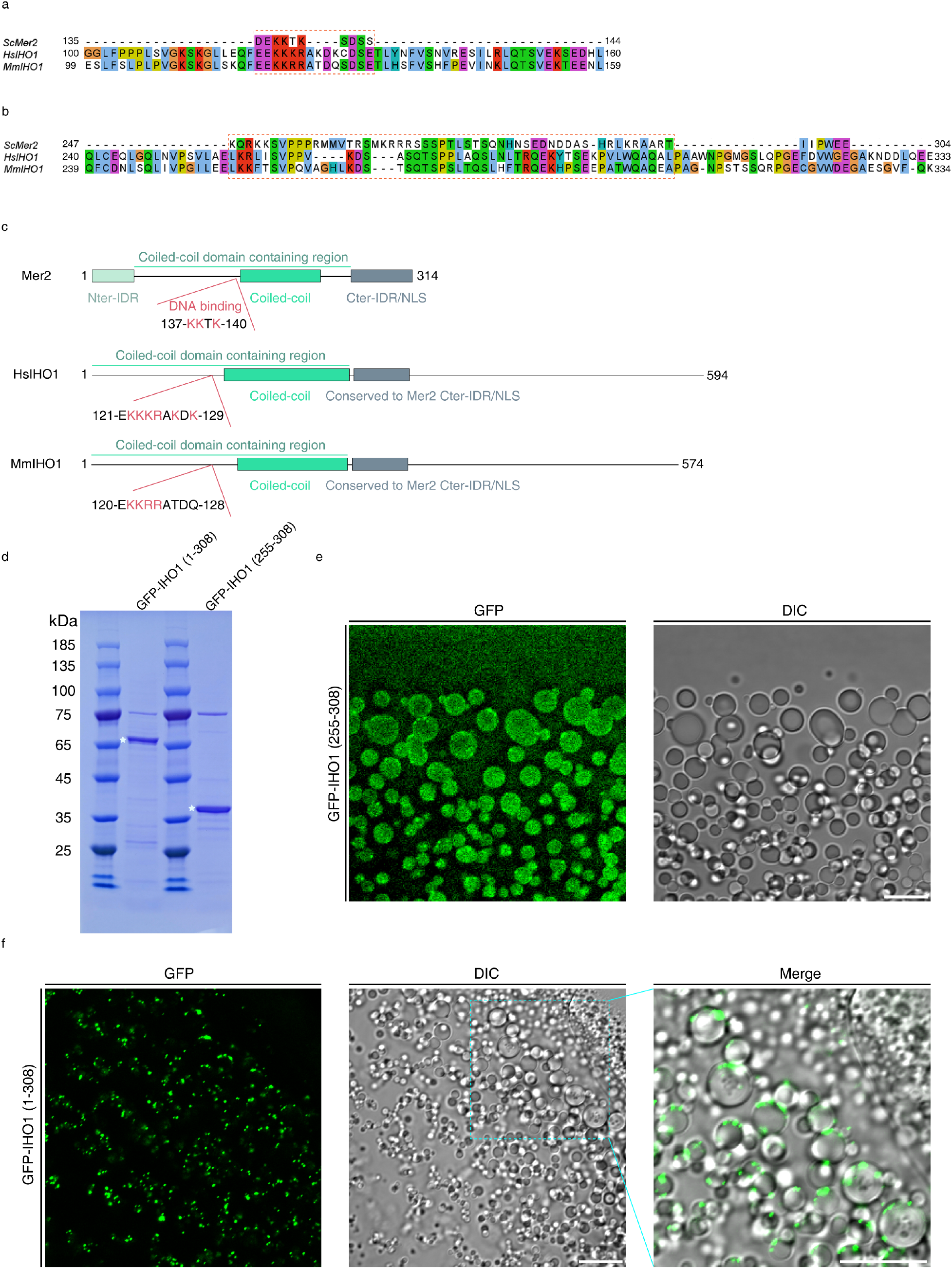
The KKTK motif and C-terminal domain of Mer2 are conserved across species. (**a**) Multiple sequence alignment of the ScMer2 KKTK motif with human and mouse Mer2 homologs using Clustal Omega. (**b**) Multiple sequence alignment of the ScMer2 C-terminal domain with human and mouse Mer2 homologs using Clustal Omega. (**c**) Domain organization of ScMer2, HsIHO1 and MmIHO1. (**d**) SDS-PGAE of GFP-IHO1 (1-308) and GFP-IHO1 (255-308). Approximately 5 μg of each protein was loaded onto a 4-12% SurePAGE gradient gel (Genscript) running in MOPS/SDS buffer and stained with Coomassie brilliant blue. Asterisks indicate proteins of interest. (**e**) Representative micrographs of liquid droplets formed by GFP-IHO1 (255-308). GFP-IHO1 (255-308) (25% PEG-3350, 9 μM) was imaged at room temperature. Scale bar, 10 μm. (**f**) Representative micrographs of liquid droplets formed by GFP-IHO1 (1-308). GFP-IHO1 (1-308) (25% PEG-3350, 5 μM) was imaged at room temperature. Unlike GFP-IHO1 (255-308), the GFP signal of GFP-IHO1 (1-308) is located on the surface of liquid droplets, whereas liquid droplets are not completely fluorescent. PEG-3350 was though to serve as a crowding agent, but it might interact with GFP-IHO1 (1-308) to form liquid droplets. Thus, GFP-IHO1 (1-308) further condenses into bright puncta on the surface of liquid droplets. Scale bars, 10 μm.

**Extended Data Fig. 3.**
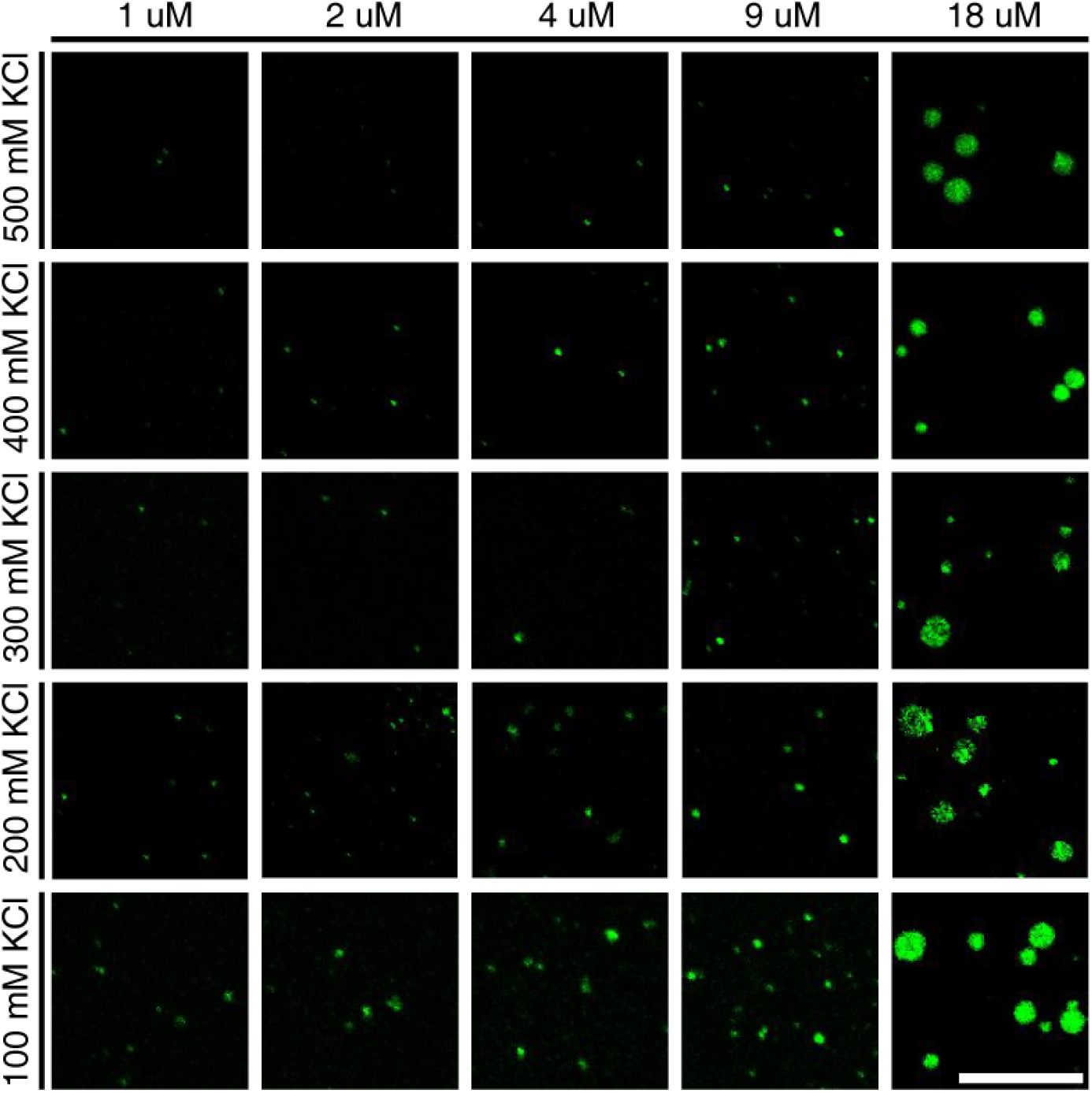
Phase diagram of Mer2. Phase diagram of Mer2 phase separation based on varying Mer2 and KCl concentrations. Scale bar, 10 μm.

**Extended Data Fig. 4.**
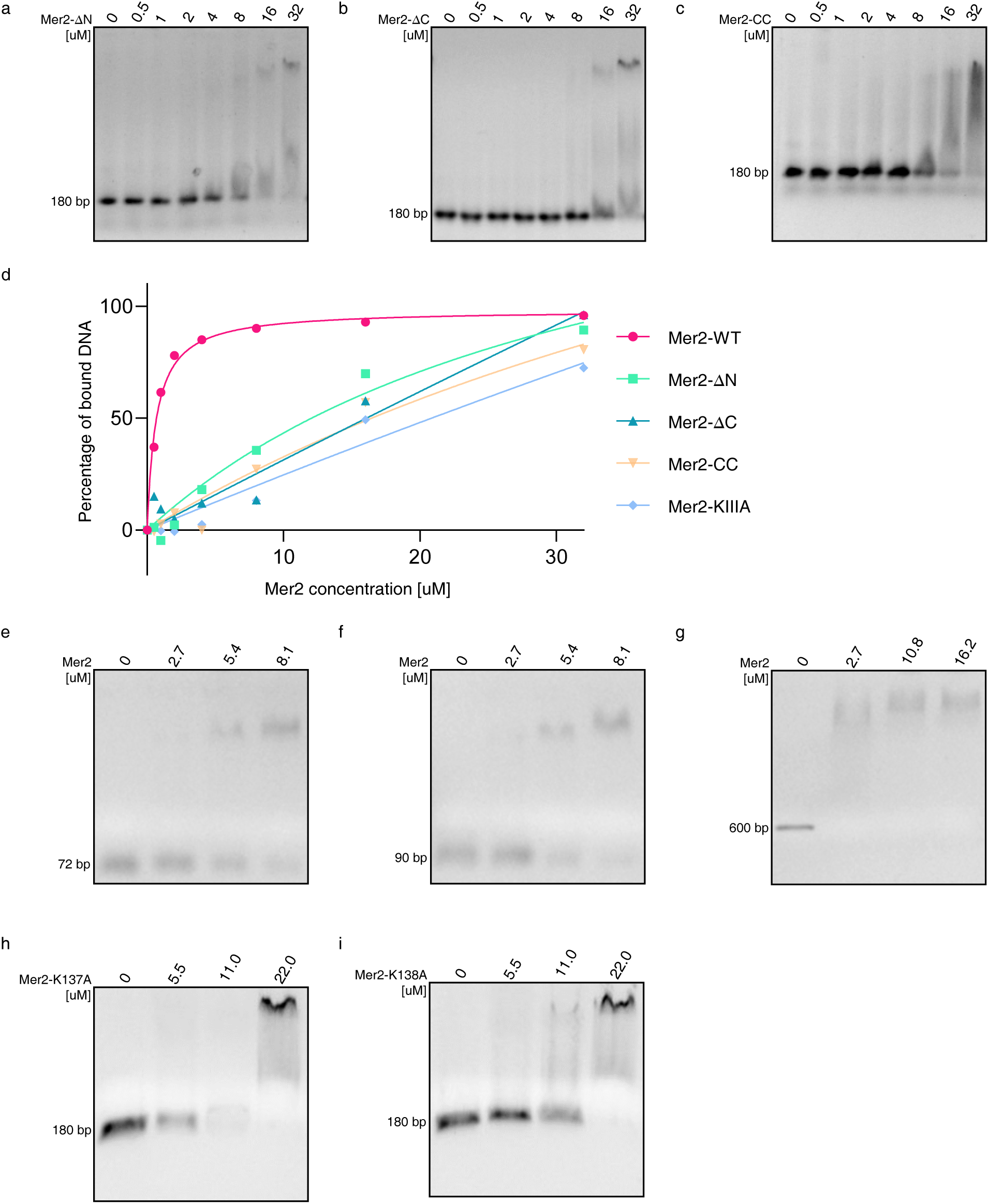
DNA binding ability of Mer2 variants. (**a**), (**b**) and (**c**) Gel-shift assay using varying concentrations of wild-type Mer2 to shift 0.1 μM 180 bp DNA in high salt buffer (500 mM KCl). (**d**) Quantification of gel-shift assays of each Mer2 variant from *n* = 3 independent experiments. (**e**), (**f**) and (**g**) Gel-shift assays of Mer2 using varying concentrations of Mer2 to shift DNA substrates that vary in length and sequence. Gel-shift assays were carried out in high salt buffer (500 mM KCl), and the concentration of each DNA substrate used was 0.02 μg/μl. (**h**) and (**i**) Gel-shift assay using varying concentrations of Mer2-K137A and Mer2-K138A to shift 180 bp DNA in high salt buffer (500 mM KCl). The concentration of DNA substrate used was 0.02 μg/μl.

**Extended Data Fig. 5.**
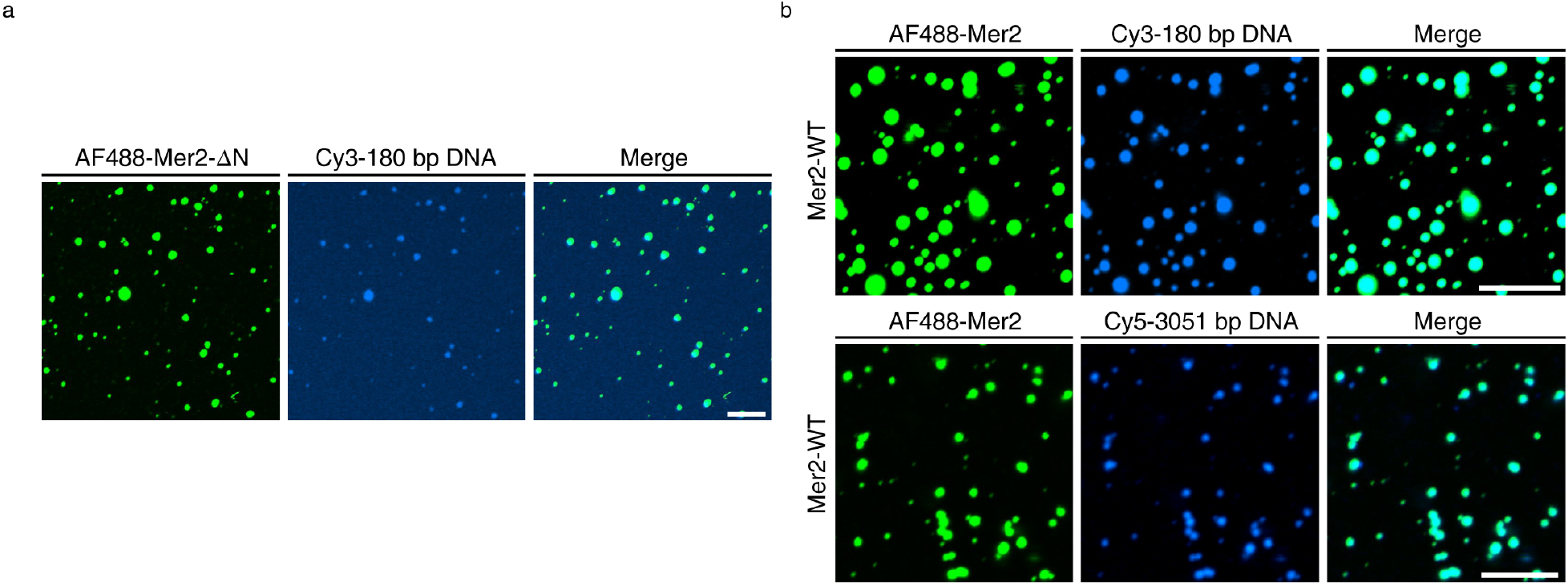
Mer2-ΔN -DNA phase separation and Mer2-DNA phase separation using different DNA molecules. (**a**) Representative micrographs of Mer2-ΔN phase separation with DNA. Mer2-ΔN (2% was labeled with Alexa Fluor 488, 10 μM) was transferred to room temperature, and 5’-Cy3-180 bp DNA (0.27 μM) was added and imaged. Scale bar, 10 μm. (**b**) Representative micrographs of Mer2-DNA phase separation. Mer2 (2% was labeled with Alexa Fluor 488, 10 μM) was transferred to room temperature, and 5’ Cy3-180 bp DNA (0.27 μM) and 5’ Cy5-3051 bp DNA (0.016 μM) were added and imaged. Scale bar, 10 μm.

**Extended Data Fig. 6.**
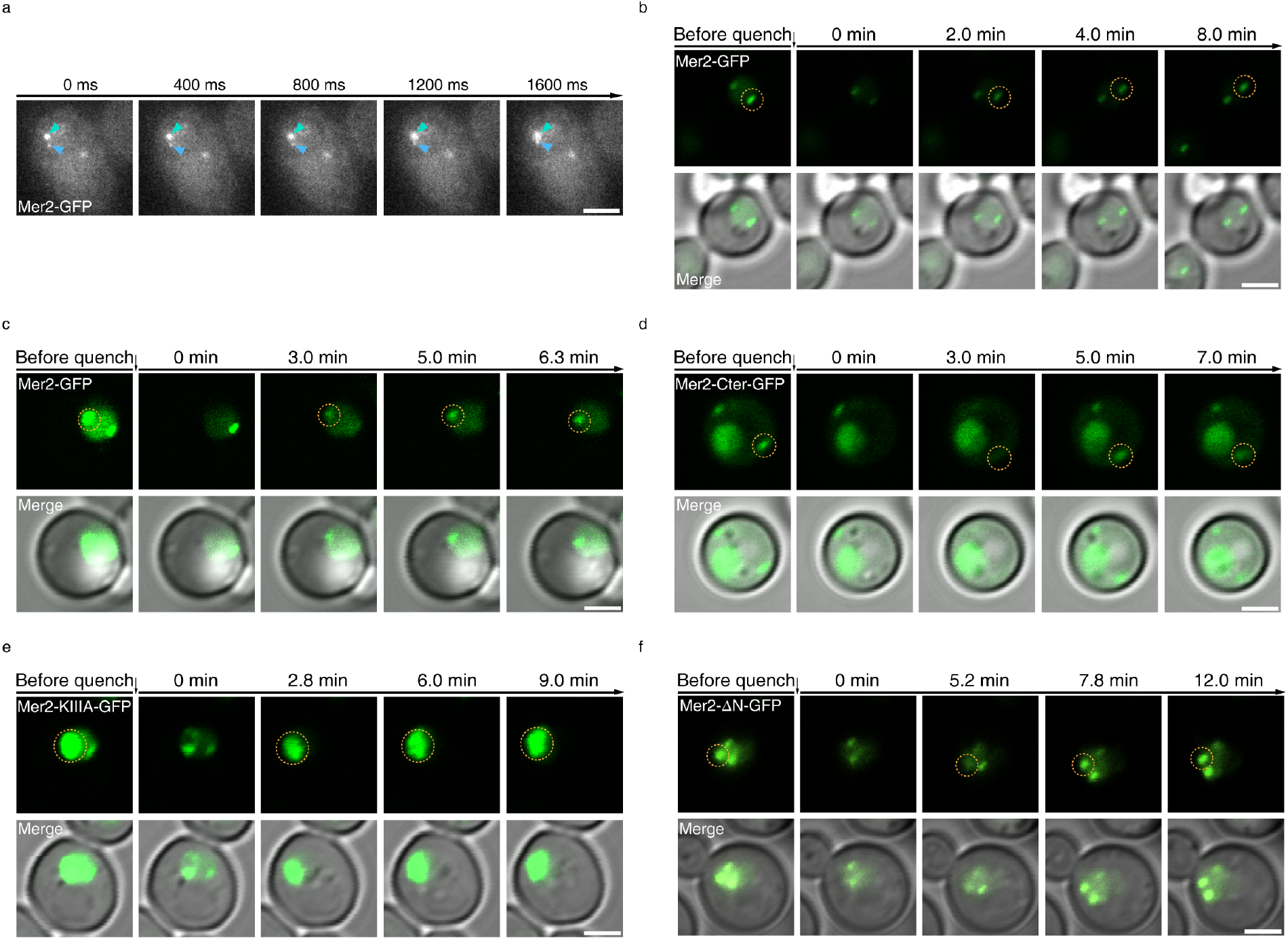
Live-cell imaging of Mer2 phase separation and FRAP analysis of Mer2 variants. (**a**) Live-cell imaging of fusion of Mer2-GFP puncta in nuclei. Yeast cells were imaged after transfer to sporulation medium for 4 hours. Green and blue arrows indicate two separate Mer2-GFP puncta that are about to fuse. Scale bars, 2 μm. (**b**) and (**c**) Representative micrographs of FRAP analysis of Mer2-GFP before and after quenching. Scale bar, 2 μm. (**d**), (**e**) and (**f**) Representative micrographs of FRAP analysis of GFP-tagged Mer2 variants before and after quenching. Scale bar, 2 μm.

**Extended Data Fig. 7.**
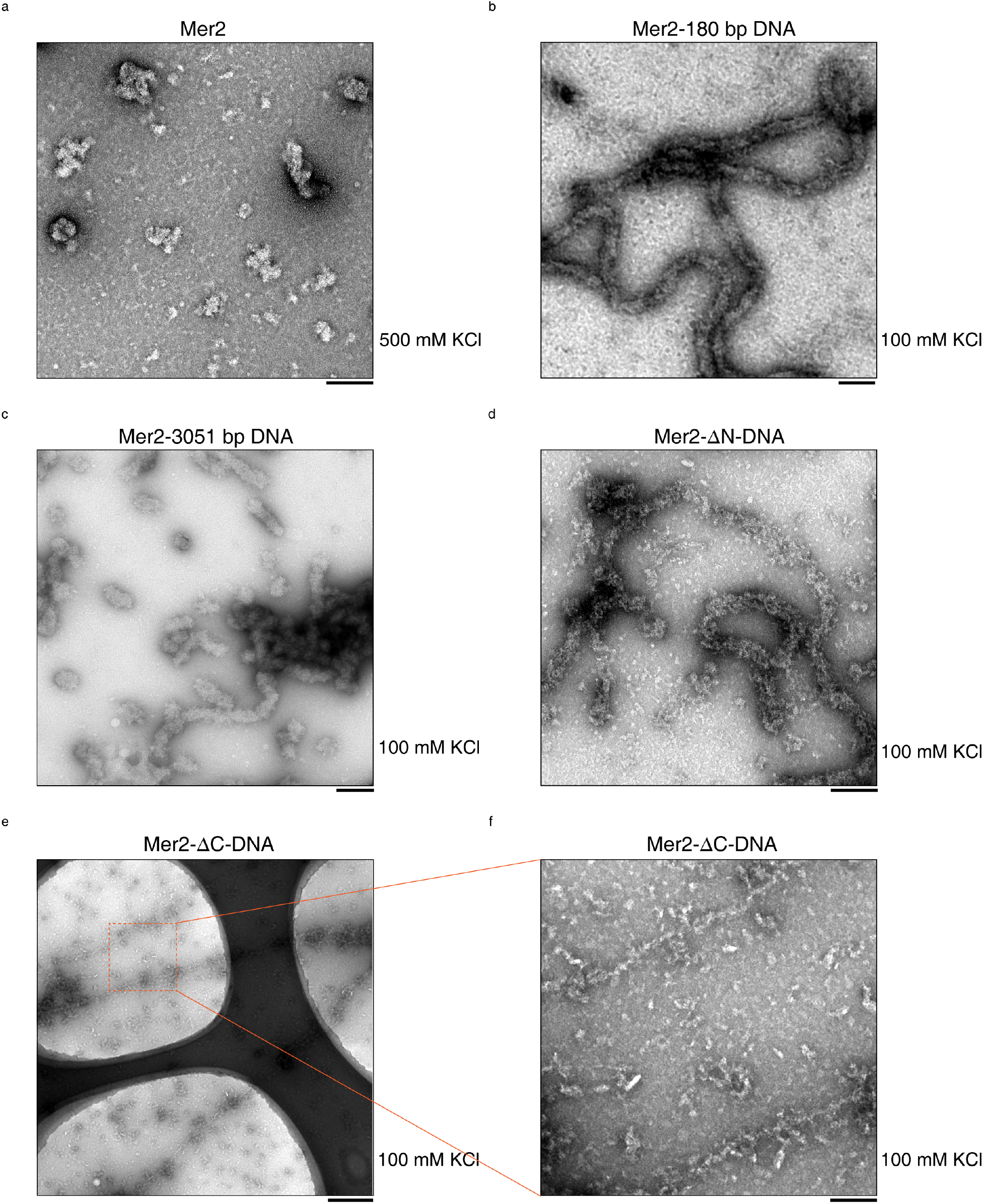
Negative-staining electron microscopy of Mer2 and Mer2-DNA filaments. (**a**) Representative negative-staining EM image of Mer2 in high salt buffer (500 mM KCl), which displayed roughly globular structures and tended to aggregate. Scale bar, 100 nm. (**b**) and (**c**) Representative negative-staining EM images of Mer2-DNA filaments in low salt buffer (100 mM KCl) using 180 bp DNA and 3051 bp DNA, both of which exhibit thick and long filamentous structures. Scale bars, 100 nm. (**d**) Representative negative-staining EM image of Mer2-ΔN-DNA filament in low salt buffer (100 mM KCl), which displayed similar structures as Mer2-DNA filament, but the Mer2-ΔN-DNA filament seemed to be less stable and tended to collapse. Scale bar, 100 nm. (**e**) Representative negative-staining EM image of the Mer2-ΔC-DNA filament at low magnification, which displayed long and thin filamentous structures. Scale bar, 500 nm. (**f**) Representative negative-staining EM image of the Mer2-ΔC-DNA filament at higher magnification, which displayed much thinner filamentous structures compared to the thick filaments formed by Mer2-DNA. Scale bar, 100 nm.

**Extended Data Fig. 8.**
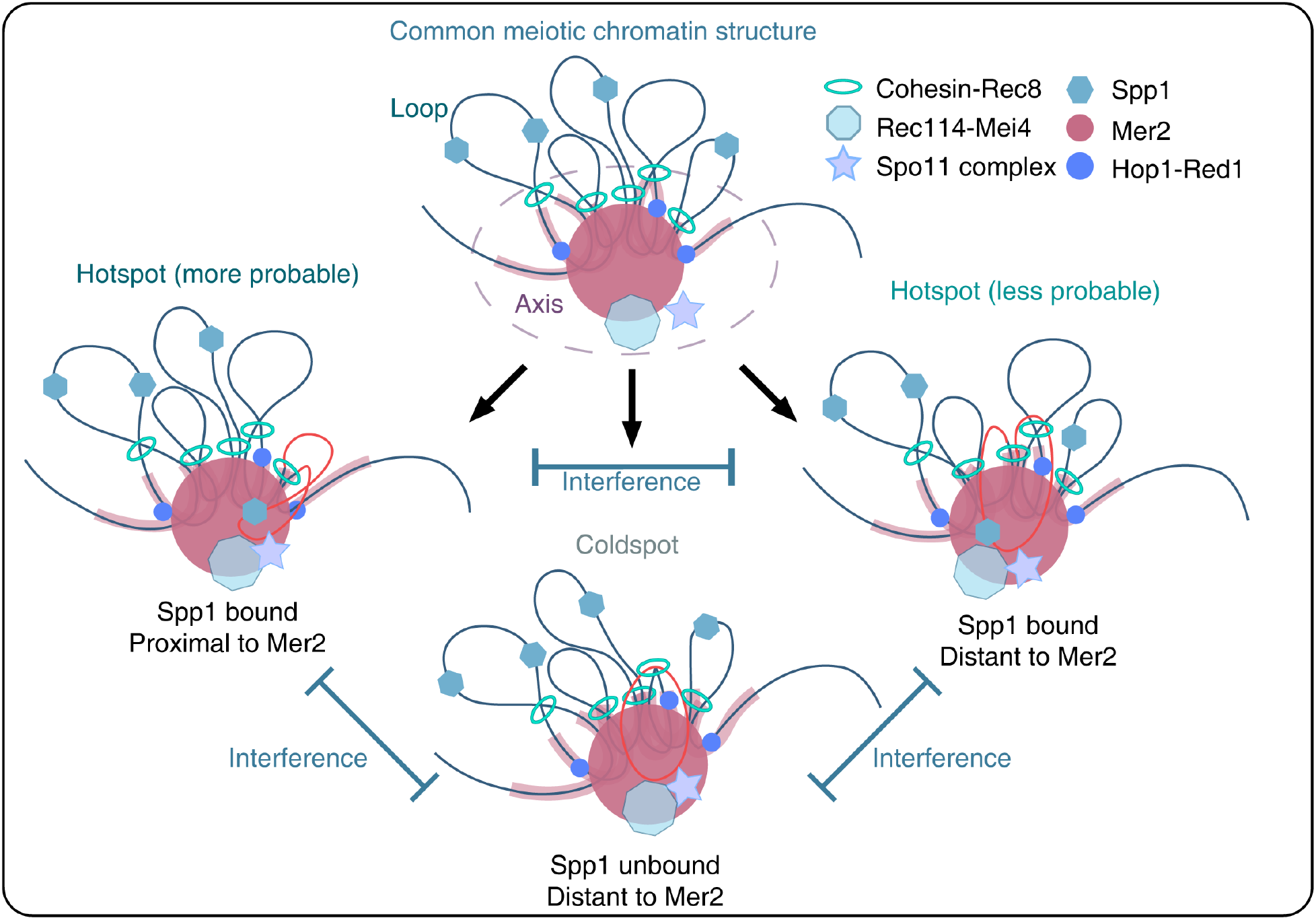
Proposed model of how phase separation of Mer2 organizes droplet-loop structure to drive and diversify programmed DSB formation during meiosis I. Phase separation of Mer2 contributes to the establishment of a common meiotic chromatin structure and organizes a droplet-loop structure, which was also characterized as a loop-axis structure. All chromatin loops within the local group can be selected for DSB formation based on whether they can be tethered to the Mer2 droplet via phase separation of Mer2-Spp1-DNA. The thermodynamics of biomolecular condensates formed via phase separation and local chromatin dynamics introduce a certain degree of randomness to loop tethering, which drives and diversifies the progress of programmed DSB formation. The competition for the limited surface area of the Mer2 droplet and other factors results in interference among spatially neighboring programmed DSB sites.

**Supplementary Video 1. Time-lapse imaging of Mer2 phase separation.** Mer2 phase separation was induced by incubating wild-type Mer2 (2% was labeled with Alexa Fluor 647, 10 μM) at 19°C and matured over 40 min. Left: bright field; middle: AF647-Mer2; right: merge of AF647-Mer2 with bright field. Scale bar, 20 μm.

**Supplementary Video 2. Time-lapse imaging of Mer2-Spp1 phase separation.** Mer2-Spp1 phase separation was induced by mixing an equal volume of wild-type Mer2 (2% was labeled with Alexa Fluor 647, 10 μM) with Spp1 (2% was labeled with Alexa Fluor 488, 10 μM), and the sample was incubated at 19°C for 20 min. Left: AF594-Mer2; middle: AF488-Spp1; right: merge of AF594-Mer2 with AF488-Spp1. Scale bar, 20 μm.

**Supplementary Video 3. Time-lapse imaging of Mer2-ΔN phase separation.** Mer2-ΔN phase separation was induced by incubating Mer2-ΔN (2% was labeled with Alexa Fluor 488, 10 μM) at 19°C over 15 min. Left: bright field; middle: AF488-Mer2-ΔN; right: merge of AF488-Mer2-ΔN with bright field. Scale bar, 20 μm.

**Supplementary Video 4. Time-lapse imaging of Mer2-DNA phase separation using 180 bp DNA.** Wild-type Mer2 (2% was labeled with Alexa Fluor 488, 10 μM) was incubated at 19°C, and 5’ Cy3-180 bp DNA (0.27 μM) was added and imaged over 40 min. Left: AF488-Mer2; middle: 5’ Cy3-180 bp DNA; right: merge of AF488-Mer2 with 5’ Cy3-180 bp DNA. Scale bar, 20 μm.

**Supplementary Video 5. Time-lapse imaging of Mer2-DNA phase separation using 3051 bp DNA.** Wild-type Mer2 (2% was labeled with Alexa Fluor 488, 10 μM) was incubated at 19°C, and 5’ Cy5-3051 bp DNA (0.016 μM) was added and imaged over 15 min. Left: AF488-Mer2; middle: 5’ Cy5-3051 bp DNA; right: merge of AF488-Mer2 with 5’ Cy3-180 bp DNA. Scale bar, 20 μm.

**Supplementary Video 6. Time-lapse imaging of Mer2-GFP in the cytoplasm during premeiotic S phase.** Diploid yeast cells with only one allele of Mer2 tagged with GFP at its C terminus were imaged using SR-FACT after transfer to sporulation medium for 2 hours. Left, Mer2-GFP imaged using Hessian-SIM; right, a slice from ODT that corresponds to the focal plane of Hessian-SIM. Scale bar, 2 μm.

**Supplementary Video 7. Time-lapse imaging of Mer2-GFP in the nucleus during meiosis I prophase.** Diploid yeast cells that have only one allele of Mer2 tagged with GFP at its C terminus were imaged using the Hessian-SIM module from SR-FACT after transfer to sporulation medium for 4 hours. Left, a slice from ODT that corresponds to the focal plane of Hessian-SIM; right, Mer2-GFP imaged using Hessian-SIM. Scale bar, 2 μm.

**Supplementary Videos 8, 9. Time-lapse imaging of Mer2-GFP in the nucleus during meiosis I prophase.** Diploid yeast cells that have only one allele of Mer2 tagged with GFP at its C terminus were imaged using the Hessian-SIM module from SR-FACT after transfer to sporulation medium for 4 hours. Scale bar, 2 μm.

**Supplementary Video 10. FRAP analysis of Mer2-GFP in yeast nucleus.** Yeast cells transformed with pCUP1-Mer2-GFP, Mer2-GFP were expressed under the control of the CUP1 promoter. The GFP signal was quenched using a 488-nm laser. Scale bar, 2 μm.

## Notes

### Competing Interest Statement

The authors have declared no competing interest.

